# MICLEAR: Intelligent Molecular Cytology for Intraoperative Margin Assessment of Pancreatic Ductal Adenocarcinoma

**DOI:** 10.1101/2023.12.01.569675

**Authors:** Tinghe Fang, Daoning Liu, Xun Chen, Keji Zhou, Chunyi Hao, Shuhua Yue

## Abstract

Pancreatic ductal adenocarcinoma (PDAC) is a highly mortal cancer whose only potentially curative treatment is surgical resection. Intraoperative assessment of its surgical margins is vital for patient survival. Frozen-section biopsy is routinely performed for this purpose. However, its high dependence on pathologists’ experience frequently poses diagnostic discrepancies. The essential invasiveness of PDAC also causes sampling errors. This study developed an intelligent molecular cytology approach with improved diagnostic objectivity and broader sampling coverage. Our method, Multi-Instance Cytology with Learned Raman Embedding (MICLEAR), is characterized by compositional information provided by label-free Raman imaging. First, 4085 cells were brushed off from the pancreases of 41 patients and imaged using stimulated Raman scattering microscopy. Then, a contrastive learning-based cell embedding model was developed to compress each cell’s morphological and compositional information into a compact cell vector. Finally, a multi-instance learning-based diagnosis model using cell vectors was employed to predict the likelihood of a patient’s margin being positive. MICLEAR achieved 80% sensitivity, 94.1% specificity, and an AUC of 0.86 on 27 patients for validation, comprising 10 with positive margins and 17 with negative ones, within approximately 8 minutes per patient. It may hold promises for more efficient and accurate intraoperative assessment of PDAC surgical margins.

## INTRODUCTION

Pancreatic ductal adenocarcinoma (PDAC) stands as the third leading cause of cancer-related deaths, with an abysmally low 5-year relative survival rate of ∼10%.^1,2^ The only potentially curative treatment currently available is surgical resection. Assessment of surgical margin status is crucial in evaluating the adequacy of resection, as it identifies residual cancer cells that significantly impact patient outcome.^3,4^ The gold standard of margin assessment, permanent section diagnosis, requires several days for diagnosis. To obtain results during operations, frozen section diagnosis, which typically takes 30 minutes,^5^ is routinely performed for significant transection margins, such as pancreatic neck margins (PNMs).^6^ Extended resection is necessary when cancer cells are detected on the margin. Following the cessation of resection, the margin tissue is sampled for permanent section diagnosis to provide the definitive margin status report.

Despite the significance of intraoperative assessment of PDAC margins, achieving this goal reliably with frozen sections is challenging. Its diagnosis depends heavily on pathologist specialization and sampling methodology,^7,8^ probably accounting for the reported false-negative rates ranging from 4.3% to 75% among centers.^9-11^ PDAC is also shown to be one of the cancers where frozen sections have the highest discordance with the final diagnosis.^12^ These two problems are primarily attributed to sampling errors, in which lesions are not cut off or included in the prepared slide,^13,14^ as well as misinterpretation due to tissue artifacts and confusion with pancreatitis.^15-17^ Moreover, PDAC is characterized by high invasiveness and dispersed growth at the invasive front,^18-21^ aggregating the risk of sampling errors. Therefore, limitations of frozen sections in PDAC margin assessment indicate the urgent need for an alternative method with enhanced diagnostic objectivity and sampling coverage.

To meet this need, several histological substitutes have been proposed. Fluorescent-guided surgery utilizes fluorescent labeling of cancers, which involves the strict selection of biomarkers and has a relatively low specificity of ∼70%.^22-24^ Optical coherence tomography visualizes tissue structures on the cross section but lacks chemical information, making its sensitivity and specificity less than 75%.^25^ *In vivo* mass spectroscopy (MS) and *ex vivo* spontaneous Raman spectroscopy achieved over 90% accuracy by recognizing the abnormal biochemical composition of PDAC tissues.^26,27^ However, MS involves a complex analytical procedure, while spontaneous Raman spectroscopy has a narrow detection volume and needs lengthy data acquisition. Stimulated Raman scattering (SRS) microscopy is a label-free imaging technique that maps Raman scattering signals arising from vibrational modes of molecules. Images acquired convey morphological and compositional information with high speed, high spatial resolution, and high chemical sensitivity. This approach has been used to assess PDAC margins by combining with second harmonic generation microscopy.^28^ Nevertheless, the high invasiveness and dispersion of PDAC requires a large field of view on tissues, which can hardly be achieved by SRS microscopy within the short periods required for intraoperative diagnosis. In summary, current histological substitutes have drawbacks including low accuracy, complex data processing, or low detection speed, making them unsuitable for intraoperative applications.

As a counterpart to histology, cytology is commonly used for cancer screening owing to its higher cost-effectiveness and lower invasiveness. In particular, cytology adapts to the high invasiveness and dispersion of PDAC through efficient margin sampling, thus allowing for compressional and rapid analysis with a variety of cells around suspected lesions. However, conventional cytology, which only exploits cellular morphology, typically has a sensitivity lower than 60%.^29-31^ For example, in the diagnosis of pancreatic lesions, brush cytology can collect cells from narrow pancreatic ducts but has a sensitivity of only ∼50%.^32,33^ This drawback could be significantly mitigated by taking abnormal molecular features of cancer cells into account. Therefore, based on the aberrant accumulation of lipid droplets (LDs) in PDAC, as revealed by SRS microscopy,^34^ we developed a refined brush cytology for assessing PDAC margins.

To predict PDAC margin status with cellular SRS images, we developed a machine learning-assisted method named Multi-Instance Cytology with Learned Raman Embedding (MICLEAR). To construct our dataset, we imaged 4085 pancreatic cells brushed off during operations at the Raman bands corresponding to lipid and protein. Normal pancreatic cells were extracted from 7 patients without pancreatic diseases, while PDAC and margin (PNM) cell samples were obtained from 34 patients with PDAC. Then, a cell embedding model was trained with a contrastive learning (CL) framework to represent each cell with compact cell vectors. Cell embedding substitutes for the traditional feature extraction workflow, which requires costly feature engineering. Finally, with a multi-instance learning (MIL) algorithm, marker cells were identified as evidence of positive margins. Our method achieved 80% sensitivity and 94.1% specificity on 27 patients with PNM cell samples for validation, including 10 with positive PNMs and 17 with negative PNMs. The test time was approximately 8 minutes for each patient. Our method may hold great potential for accurate, easy-to-use, and rapid intraoperative assessment of PDAC margins.

## RESULTS

### Workflow of Multi-instance Cytology with Learned Raman Embedding (MICLEAR)

The workflow of our developed MICLEAR is illustrated in **Fig. 1**. Exfoliative cell samples were collected from three sources using medical cytology brushes, including healthy pancreases (*N* samples), PDAC tumor cores (*T* samples), and PNM tissues (*M* samples) (**Fig. 1a**). *N* and *T* samples corresponded to cell populations without and with cancer cells, respectively. Whether *M* samples contained cancer cells depended on pathological reports. Cellular images were acquired by two-color SRS microscopy at the Raman bands for CH_2_ stretching in lipid around 2850 cm^-1^ and for CH_3_ stretching in protein around 2930 cm^-1^, respectively (**Fig. 1b**). Example SRS images of *N* and *T* samples are shown in **Fig. S1**. Individual cells were segmented from SRS images at 2930 cm^-1^ with a pre-trained deep learning model (**Fig. 1c**). The obtained segmentation mask was applied to the corresponding SRS images at two channels to produce single-cell SRS images, with which single-cell molecular images reflecting lipid and protein concentration were calculated (**Fig. 1d**). Then, with single-cell molecular images of *N* and *T* samples, a cell embedding model was trained to transform each single-cell molecular image into a 512-dimensional cell vector, which was a compact representation of single-cell morphological and compositional information (**Fig. 1e**). Finally, a MIL-based model was trained to predict the positive probability of a sample according to all cell vectors within. This model was evaluated with leave-one-out cross-validation (LOOCV), in which one *M* sample was left out for testing, while *N, T*, and the remaining *M* samples were used for training (**Fig. 1f**). The evaluation metrics, including sensitivity, specificity, and area under the receiver operating characteristic curve (AUC), were calculated from the testing results.

**Figure 1.**
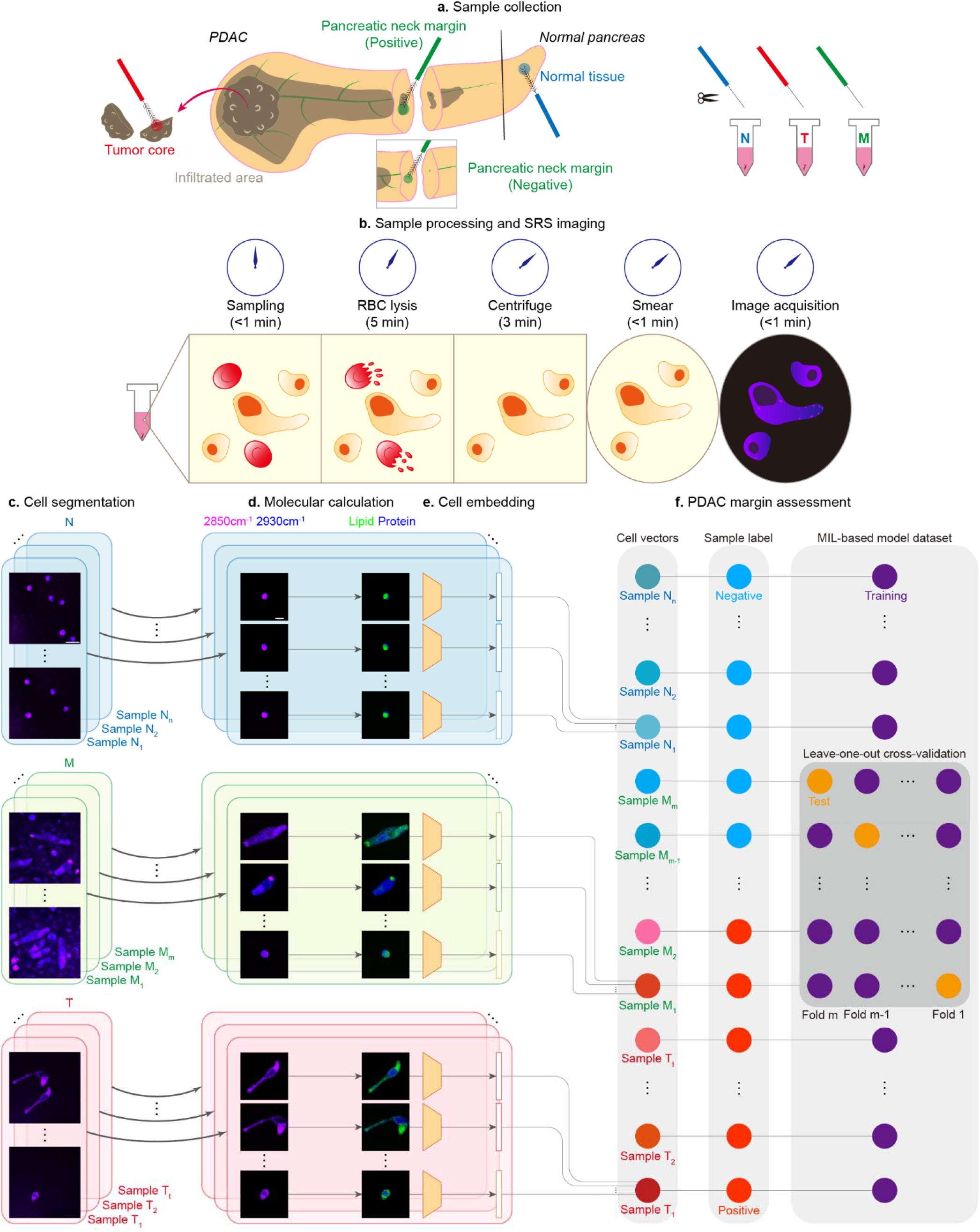
Overview of the MICLEAR-assisted molecular cytology. **a**, Exfoliated cell collection from normal pancreatic tissues (*N* samples), PDAC tumor cores (*T* samples), and PNMs (*M* samples). **b**, Sample processing and two-color SRS imaging of cells. **c**, Single-cell segmentation. **d**, Calculation of molecular images reflecting lipid and protein concentration. **e**, Cell embedding with a model trained under the CL framework. **f**, PDAC margin assessment with an MIL-based model. Scale bars, 20 μm (**c**), 10 μm (**d**).

### Single-cell Segmentation and Extraction of Compositional Information

Our home-built SRS microscope setup is shown in **Fig. 2a**. For single-cell segmentation from SRS images, we applied *Cellpose*,^35^ a well-developed deep learning algorithm based on intensity gradients. Compared with watershed, a traditional method, and *Stardist*,^36^ another deep learning-based method, *Cellpose* performed better (0.86 global Dice coefficient, 1.03 cell number ratio, and 0.86 cellwise Dice coefficient, as defined in the *Methods* section), especially for cells with nonconvex contours (**Fig. S2**). Then, the Raman spectra of pure chemical samples (**Fig. 2b**) were used to decompose two-color SRS images into signals from lipid and protein, which were proportional to their concentrations.^37^ This processing produced single-cell molecular images composed of two chemical channels. In the lipid channel, some cells in *T* samples have obviously more LDs (**Fig. 2c**). To quantify the difference between cells in *N* and *T* samples, we manually extracted twelve cell features, including five morphological and seven compositional ones, from SRS images (**Fig. S3**). Among these features, different feature value distributions between *N* and *T* samples were seen, such as in two morphological features (area, shape factor) and two compositional ones (LD number and LD dispersity) (**Fig. 2d**). For these features, *T* samples had a significant proportion of cells with saliently higher feature values than all cells in *N* samples. This implies that cancer cells may have larger area, more irregular shape, more LDs, and more dispersed LDs. The potential contribution of compositional information to accurate diagnosis can also be seen.

**Figure 2.**
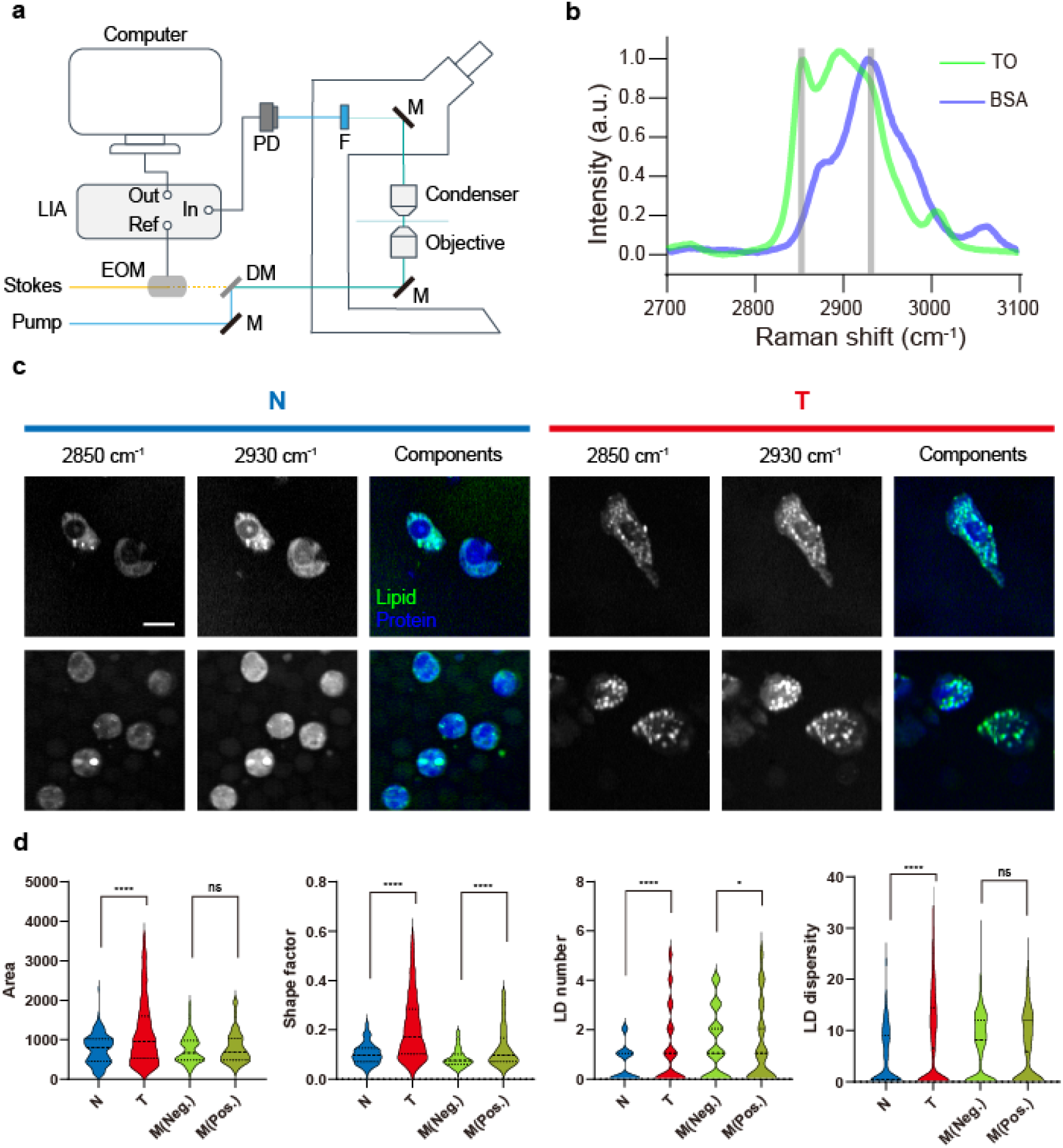
SRS image acquisition and preprocessing. **a**, Setup of the SRS microscope. DM, dichroic mirror. EOM, electro-optic modulator. F, optical filter. LIA, lock-in amplifier. M, mirror. PD, photodetector. Ref, reference signal. **b**, Raman spectra of triolein (TO) and bovine serum albumin (BSA). Gray columns denote two Raman shifts used for two-color SRS microscopy, 2850 cm^-1^ and 2930 cm^-1^. **c**, Two-color SRS images and corresponding molecular images in *N* and *T* samples. **d**, Feature distribution in *N, T, M* (negative), and *M* (positive) samples; outliers were removed as described in the *Methods* section; the area was quantified by the number of pixels with constant pixel size. Scale bars, 10 μm. ns: no statistically significant difference; *: p<0.05; ****: p<0.0001; tested by Mann-Whitney U test.

Despite different cell feature distributions in *N* and *T* samples, there will be two major problems if these features are used to predict whether a sample has cancer cells with conventional machine learning. On the one hand, whether a cell feature contributes to accurate diagnosis can only be known after feature extraction, which is costly and may be a waste of resources. On the other hand, essential overlaps of feature distributions existed between *N* and *T* samples, severely impairing the performance of models using sample-level statistics, such as average and maximal feature values. Huge feature value overlaps between *N* and *T* samples were also seen in other extracted cell features (**Fig. S4**). Furthermore, the feature value difference may weaken between *M* (Negative) and *M* (Positive) samples (**Fig. 2d, Fig. S4**), which may be attributed to a smaller proportion of cancer cells or their less significant malignancy. To address the two issues above, MICLEAR incorporated two crucial functions. One was CL-based automatic feature discovery and extraction for compact representation of cells. The other was MIL-based recognition of marker cells that indicated sample positivity.

### Contrastive Learning for Single-cell Dimensionality Reduction into Cell Vectors

To acquire compact cell representations with minimal prior knowledge about valuable features, we first developed a cell embedding model inspired by *SimCLR*,^38^ a simple CL framework. Through this process, a single-cell molecular image with 400×400 pixels was compressed into a 512-dimensional cell vector by more than 300 times, whose components denoted high-dimensional cell features and were associated with cell morphology and composition. The model training attempted to increase the difference between the output vectors of different cells and decrease the difference between the output vectors of the same cell with random rigid transformations (**Fig. 3a**). In comparison with *SimCLR* where the output vectors were fed into a linear classifier for supervised classification, we applied them to a MIL classifier for weak-supervised classification. Another difference from *SimCLR* was that *SimCLR* focused on refining model weights for optimal feature space and transferability, while ours focused more on utilizing the output vectors instead of what weights were learned by the model.

**Figure 3.**
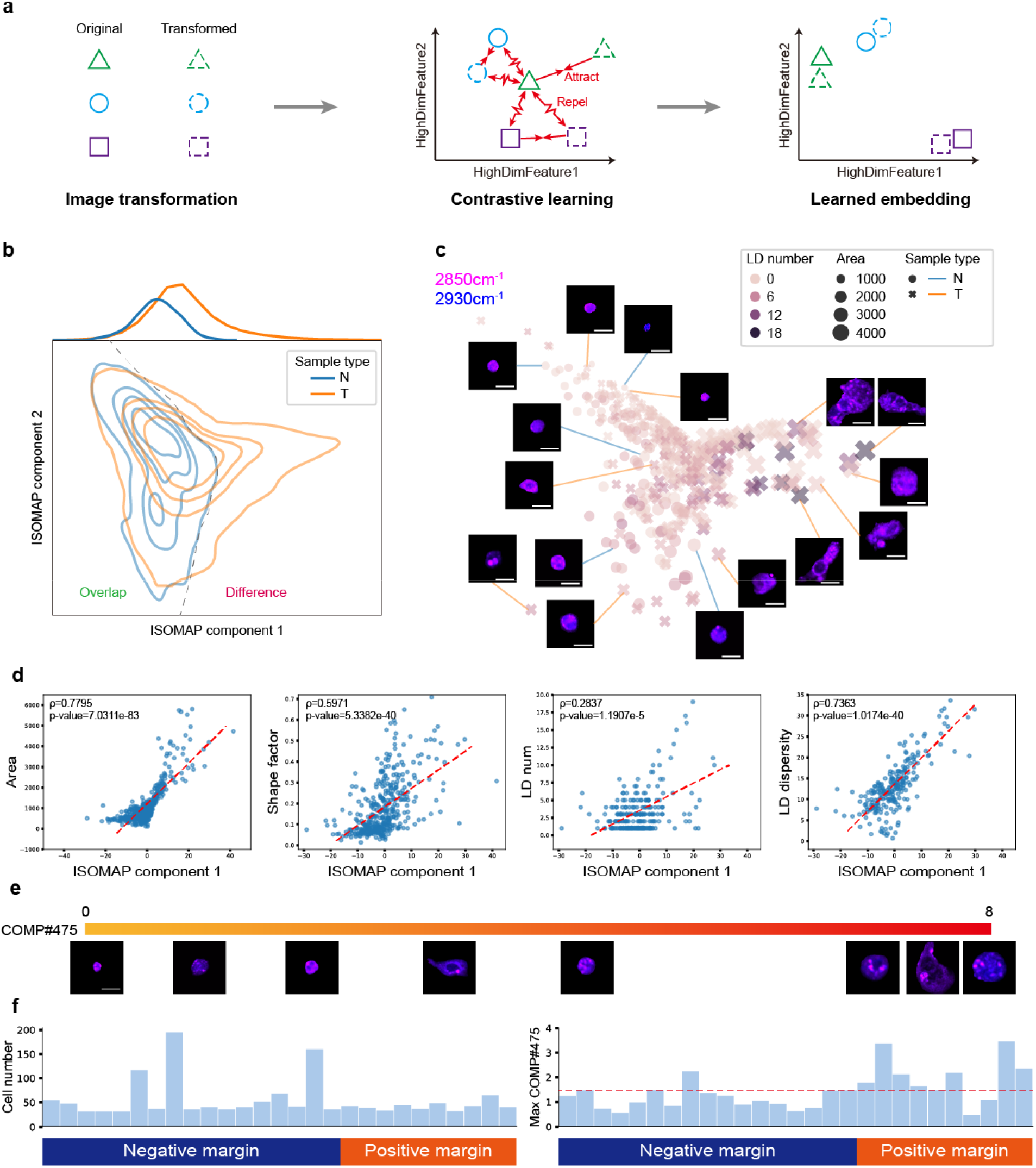
CL-based cell embedding model. **a**, Diagram of the CL framework. **b**, ISOMAP of cell vectors in *N* and *T* samples; the contour plot of kernel density estimation result is shown. **c**, ISOMAP in **b** associated with cell images and features. **d**, Correlation between cell features and the first component of ISOMAP, tested by Spearman’s rank correlation test. Cells without LDs were excluded from the correlation analysis between ISOMAP component 1 and two LD-related features. Red dashed lines denote linear regression between values of ISOMAP component 1 and cell features. Cells are randomly down-sampled by a factor of 10 for visual clarity in **c** and **d. e**, Representative cell images with their COMP#475 values. **f**, Cell numbers of *M* samples and the COMP#475 maximum among their cells. The red dashed line denotes an approximate boundary of the COMP#475 maximum between negative and positive margins. Scale bars, 10 μm.

After training the cell embedding model, we analyzed the cell vectors’ distribution and correlation with cell features. ISOMAP,^39^ a manifold learning method that maintains the geodesic distance between points, was used to reduce the dimensionality of cell vectors into two to visualize vector distribution (**Fig. 3b**). A key finding was that cell vectors in *N* samples basically constituted a subset of those in the *T* samples. Therefore, we hypothesized that the difference area between *N* and *T* samples consisted of cells that indicated positive samples. These cells generally had larger areas, more irregular shapes, more LDs, and/or more dispersed LDs (**Fig. 3c**). Spearman’s rank correlation tests showed Spearman’s rank correlation coefficients (SCCs) larger than 0.5 between the first ISOMAP component and three cell features, including area, shape factor, and LD dispersity (**Fig. 3d**). These results suggest the cell vectors obtained by our cell embedding model were meaningful representations that correlated well with cell features and may facilitate the following MIL-based model for PDAC margin assessment.

We also analyzed the relationship between cell malignancy and cell vector component values. For this purpose, a vector component with a high coefficient of variation (CV) and high SCCs with cell features was preferable. The former meant that the component reflected intercellular differences well, and the latter is beneficial to correlate the component with the degree of malignancy. Here, the cell vector component COMP#475 was selected, which had top 25% CV as well as SCCs with cell area and LD dispersity larger than 0.65. On the cell level, cells with larger COMP#475 values appeared more obvious features that may correlate with malignancies, such as larger area, more irregular shape, and more LDs (**Fig. 3e**). On the patient level, except POS#7 and POS#8, the maximal COMP#475 values among all cell vectors within a sample were larger in *M* (Positive) samples than in *M* (Negative) ones (**Fig. 3f**). For other vector components with high CV, higher component values similarly occurred in most *M* (Positive) samples (**Fig. S5**). In contrast, such difference did not occur in mean values of cell features (**Fig. S6**). These results indicate that cell vectors well reflect the difference between positive and negative PDAC margins, providing the basis for margin assessment with them.

### Multi-Instance Learning Using Cell Vectors for Accurate PDAC Margin Assessment

To utilize cell vectors for PDAC margin assessment, we trained a MIL-based logistic regression (LR) model, named MI-LR. An LR model assigns a coefficient for each vector component to quantify its relevance in a positive diagnosis. Inspired by a MIL algorithm based on support vector machine (SVM), MI-SVM,^40^ our MI-LR model can predict both on the cell level and on the patient level. During training, cell-level pseudo labels assigned by the model were iteratively updated until the model converged. After training, the MI-LR model separated cells into two subgroups, significant and insignificant cells, with a positive probability threshold of 50%. The existence of significant cells in a sample meant that the sample was positive. Significant cells in *M* (Positive) samples showed distinct appearances from those of insignificant ones (**Fig. S7**). Higher positive probability was concurrent with changes in cell area, cell shape, and LDs (**Fig. S8**), suggesting that the positive probability was not predominantly determined by any one of cell features, but by a combination of them. The maximal positive probability of all cells collected from the patient was termed the patient-level positive probability and used to predict whether the patients had a positive PDAC margin. The diagnosis model achieved 80% sensitivity, 94.1% specificity, 88.9% accuracy, and an AUC of 0.86 on 27 patients with *M* samples (**Fig. 4a**). For clarity, because of difficulty in acquiring single-cell ground truths, we did not attempt to evaluate the accuracy of cell-level classification, i.e., whether significant cells were equivalent to cancer cells. Instead, we validated the contribution of the identified subgroup of significant cells to accurate diagnoses on the patient level.

**Figure 4.**
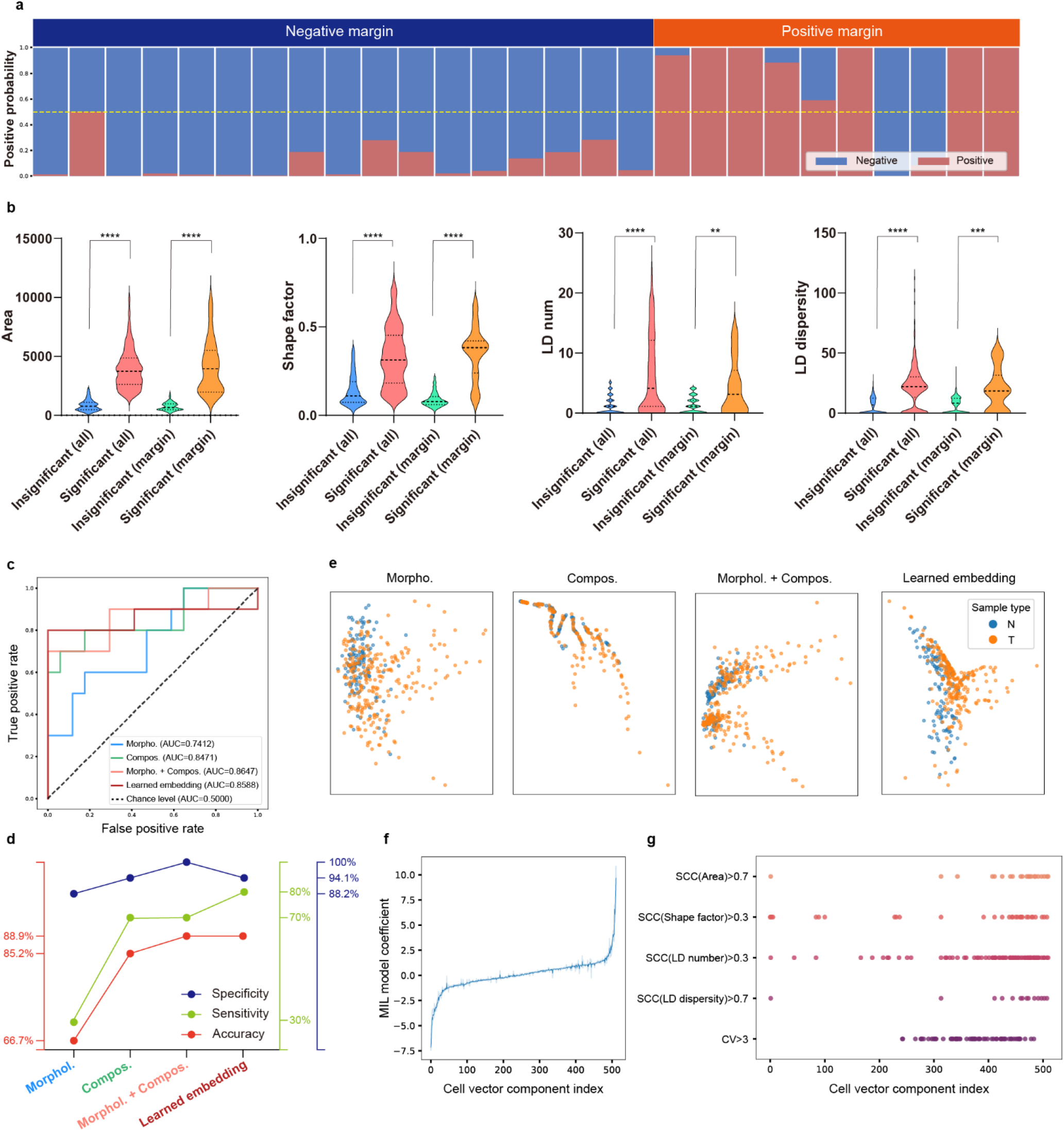
MIL-based PDAC margin assessment model. **a**, Predicted positive probability of *M* samples. **b**, Cell feature distribution of significant and insignificant cells. The feature value ranges are different from those in **Fig. 2d** because of different removed outliers. **c**, ROC curves and AUC of PDAC margin assessment with different feature sets. Morphol, extracted morphological features. Compos., extracted compositional features. **d**, Specificity, sensitivity, and accuracy of PDAC margin assessment with different feature sets. **e**, ISOMAP of different feature sets. Cells are randomly down-sampled by a factor of 10 for visual clarity. **f**, MI-LR model coefficients sorted by their mean values among all LOOCV folds in an ascending order. Light blue shows the range of each coefficient and dark blue shows the median. **g**, Cell vector components with high SCCs with cell features and high CV. **: p<0.01; ***: p<0.001; ****: p<0.0001; tested by Mann-Whitney U test.

Moreover, we extracted the distributions of previously mentioned four cell features (area, shape factor, LD number, and LD dispersity) of significant and insignificant cells (**Fig. 4b**) in *N* and *T* samples and compared them with the gross feature distributions in *N* and *T* samples (**Fig. 2d**). Between two cell subgroups separated by our MIL-based model, overlap of feature values was substantially smaller. Such reduction of feature overlaps can be similarly seen in *M* (positive) and *M* (negative) samples. Besides, even after removing the outliers for clear graphs, cells with obviously abnormal area, shape, and LDs were still included (**Fig. 4b**). However, without the MIL-based model, these cells were overwhelmed by those similar to *N* sample cells and regarded as removable outliers (**Fig. 2d**). These findings show the contribution of the MIL-based model’s ability to recognize significant cells in the sample and thus increase the accuracy of PDAC margin assessment. We also compared other cell features before and after significant cell recognition by the MIL-based model (**Fig. S4**), in which enhanced difference between two subgroups was also seen in cell perimeter and cytoplasm area. Interestingly, the difference in some cell features weakened between two subgroups, which may be because the difference in these features between *M* (positive) and *M* (negative) samples was occasional, so they were not reliable features to recognize abnormal cells.

### Interpretation of Feature Contributions in PDAC Margin Assessment by MICLEAR

The compositional information obtained by SRS microscopy is an essential improvement of our molecular cytology compared with the traditional one, which only utilizes cell morphology. To demonstrate the significance of introducing compositional information in PDAC margin assessment, we analyzed how the performance of the MIL-based model decreased when only manually extracted morphological or compositional features were used. To find out whether our cell embedding model gave out comparable cellular information to manually extracted cell features, we also compared the performance of the MIL-based model using cell vectors and using cell features.

When cell features were used for PDAC margin assessment, the MIL-based model performed better when combining morphological and compositional features than using either of them (0.86 AUC with all cell features, 0.74 AUC with morphological ones, and 0.85 AUC with compositional ones) (**Fig. 4c**). When compositional features were supplemented to morphological ones, the sensitivity increased substantially from 30% to 80%. Similarly, when morphological features were supplemented to compositional ones, the specificity and accuracy increased from 94.1% to 100% and from 85.2% to 88.9%, respectively (**Fig. 4d**). ISOMAP reflected that distribution of cell vectors and of all cell features were more dispersed than that of either feature set (**Fig. 4e**), which may be beneficial for the MIL-based model to identify significant cells. The MIL-based model using cell vectors showed similar performance to that using all cell features. Despite this, the former still holds a significant advantage because it does not need task-specific prior knowledge, which is costly to accumulate through enormous statistical analyses and experience of feature engineers. Besides, designing robust feature extraction algorithms that apply to extensive image datasets with is challenging.

Moreover, we analyzed the contribution of each cell vector component to positive diagnosis, which was reflected by the MIL-based model coefficients. They changed negligibly among different folds of the LOOCV, so could be represented with their mean values among all folds (**Fig. 4f**). We sorted cell vector components according to their MIL coefficients in an ascending order. In this way, components after COMP#245 had positive MIL coefficients, suggesting a positive correlation with cell malignancy. These components generally correlated better with malignancy-related cell features (such as larger cell area, more irregular cell shape, and more disperse LDs) and/or had higher CVs (**Fig. 4g**). For example, COMP#462 had SCCs with cell area and LD dispersity larger than 0.7; COMP#478 had a CV larger than 3, which was larger than most cell vector components, and SCCs with area and LD dispersity larger than 0.7. These results suggest that the MIL-based model may assign most weights to the cell vector components that correlated better with malignancy-related cell features and that varied greatly among cells. The latter reflects that the outcome of the CL framework, which was to produce cell vectors that distinguish cells as much as possible, was well exploited by the MIL-based model. MIL-based model coefficients, CVs, and SCCs of cell vector components are listed in **Table S1**.

### Comparison with Other PDAC Margin Assessment Algorithms Using Cell Vectors

To further verify the significance of using MIL algorithm for PDAC margin assessment, we compared our MI-LR’s performance with two categories of non-MIL algorithms. Cell vectors produced by the same cell embedding model were their inputs. One category of algorithms utilized a conventional LR model and patient-level statistics calculated with cell vectors (**Fig. S9a**). When the mean value of all cell vectors belonging to each patient was used, the sensitivity declined sharply from 80% to 50% because the malignancy-related characteristics were seriously diluted. When the maximum of each cell vector component was used, the specificity declined from 94.1% to 82.4% because this was equivalent to the situation that there was a cell incorporating the malignancy-related characteristics of all cells in the sample, which may cause false positives. Supplementing the minimum of each cell vector component to the maximum did not help. In summary, these methods have inherent limitations due to their inability to recognize a significant subgroup of cells and focus on them for PDAC margin assessment.

The other category of algorithms calculated patient-level statistics within a cluster of cells instead of within all of them. K-Means clustering was performed after dimensionality reduction via principal component analysis (PCA) or ISOMAP. The range of the selected cell cluster was similar to the difference area in **Fig. 3b**. This method was similar to K-PCA used in our previous study,^41^ but different in that the dimensionality reduction and clustering were performed on all *N* and *T* samples and then the learned model parameters were applied to each *M* sample instead of learning parameters directly on each *M* sample because exfoliated pancreatic cells brushed off from pancreatic margins were less than those collected from ascetic fluid.^41^ The methods used here were named K-N-PCA or K-N-ISOMAP for distinction (**Fig. S9b**). K-N-ISOMAP worked better than K-N-PCA (sensitivity of 80% vs 60%, specificity of 88.2% vs 82.4%, and accuracy of 81.5% vs 77.8%) but worse than MI-LR (specificity of 88.2% vs 94.1% and accuracy of 81.5% vs 88.9%). To note, K-N-PCA and K-N-ISOMAP implicitly utilized the prior knowledge that significant cells were centered in a specific cluster and could be identified with clustering, which is not so obvious. In comparison, although a special cluster (the difference area in **Fig. 3b**) was indeed found in our analysis of cell embedding results, MI-LR did not require such prior knowledge at all and, thus, may be more generalizable. Besides, after K-Means clustering, cluster to be used for margin assessment needs to be manually selected, which was not required by MI-LR.

Moreover, we compared the performance of different MIL-based model training schemes and different base machine learning algorithms of the MIL-based model (**Fig. S10**). MI- and mi-models in the literature have similar training schemes that require iterative training to make the model converge.^40^ Their difference is that for cells in each positive sample, mi-models use all of them for the next training iteration, while MI-ones only use the cell with the highest positive probability. Therefore, MI-models conform more to the fact that cells whose malignant degrees are not the highest do not make significant difference to the patient-level diagnosis, while mi-ones could regard too many cells as significant ones and cause more false positives. Replacing MI-models with mi-ones decreased the specificity from 100% to 64.7% for SVM and 94.1% to 70.6% for LR. Among MI-models, though MI-SVM had a slightly higher specificity than MI-LR (100% vs 94.1%), MI-LR reached a much higher sensitivity (80% vs 50%). Collectively, MI-LR performed better than three other MIL-based models: MI-SVM, mi-SVM, and mi-LR.

## DISCUSSION

In this study, we developed a molecular cytology based on label-free SBS microscopy and the MICLEAR algorithm developed by us for intraoperative PDAC margin assessment. Two-color SRS microscopy was used to obtain single-cell morphology and compositional information of lipid and protein. CL-based cell embedding condensed the spatially-resolved cellular Raman profiles into compact cell vectors that aimed to distinguish cells as much as possible without losing key diagnostic features. Then, an MIL-based model was used to provide patient-level positive probability with cell vectors belonging to the patient. The significance of our study to intelligent intraoperative PDAC margin assessment with high objectivity and sampling efficiency is discussed below.

By incorporating compositional information inaccessible to conventional cytology, we achieved PDAC margin assessment with comparable accuracy to routine frozen sections. In our study, comparison between different feature sets reflects the predominant contribution of compositional information to the sensitivity of PDAC margin assessment. Similar effects have been shown in spontaneous Raman spectroscopy or SRS microscopy.^41,42^ To be more specific, our study demonstrates how LD accumulation contributes to PDAC margin assessment, complying with abnormal lipid metabolism of PDAC cells reported in the literature.^34^ The substantially enhanced detection sensitivity by introducing compositional information was also reported in our previous study on the detection of peritoneally metastatic gastric cancer cells in ascetic fluid.^41^ From a biological perspective, altered metabolism has been found to occur in the early phase of cancer before significant morphological changes, accounting for the sensitivity it adds to diagnosis.^43^

Our method may have the potential to overcome the inherent limitations of frozen section-based intraoperative PDAC margin assessment by easily extending the scope of assessed margins. Some of previous studies reported that intraoperative extended resection according to frozen section diagnosis did not improve the prognosis significantly.^44^ Besides being affected by challenges in accurate and consistent pancreatic histopathology, effectiveness of frozen sections may be restrained by the limited number of margins to be assessed with standardization, especially the absence of superior mesenteric artery (SMA) margins, which have a high positivity rate (e.g. 35%)^45^ when the routinely assessed PNM is positive. Despite ambiguity in the definition of SMA margin range in histology, our cytology-based method allows direct detection of residual cancer cells over a broad range on the SMA. Accurate intraoperative assessment of SMA margins could enable more active treatment by surgeons, such as the intraoperative radiotherapy therapy.

The cell embedding model minimizes prior knowledge required for diagnosis. With customized architectures and loss functions, neural networks can automatically compress high-dimensional image data into low-dimensional feature vectors. A CL framework can embed data while optimizing the difference between output vectors. It has been widely applied in medical data to facilitate multimodal fusion and downstream diagnosis. For example, weakly-supervised CL can adjust similarities between low-dimensional representations of image patches according to patient-level labels,^46^ while self-supervised CL requires no data labels.^47,48^ Considering high intercellular heterogeneity in positive exfoliated cell samples, MICLEAR adapted the idea of self-supervised *SimCLR* for cell embedding. In this way, training of the cell embedding model did not need labeling of any cell. Otherwise, if all cells in *T* and *M* (positive) samples were assigned positive labels indiscriminately, huge label noise may be incurred.

Our scheme of MIL using cell vectors provides single-cell interpretability. Previous artificial intelligence-assisted digital histology and cytology using MIL typically comprised image patch embedding, data aggregation, and prediction.^49,50^ The diagnosis results were interpreted with attention mechanisms or saliency maps.^51,52^ These methods attribute model prediction to particular regions of image patches, which cannot provide clear cell-level interpretation and may be inaccurate because high attention is not an agent of high significance in diagnosis. In comparison, with single-cell segmentation and embedding, MICLEAR directly recognizes marker cells indicating positive samples and allows comparison between the features of them and other cells. As shown in **Fig. 4d** and **Fig. S4**, in addition to larger size and nucleocytoplasmic ratio in morphology, the lipid alteration-associated features of significant cells were revealed, including increasing and more dispersive LDs and higher lipid intensity within the cytoplasm. The nice interpretability of our method can be attributed to a designed combination of deep learning (i.e., CL) and traditional machine learning (i.e., MI-LR) instead of an end-to-end deep learning, which may impede clear interpretation a lot.

We investigated two false negatives (POS#7, POS#8) and one false positive (NEG#2) in PDAC margin assessment. Two false negatives had cell vector components similar to those of *M* (Negative) samples (**Fig. S5**). They did not have cells similar to the significant cells in other *M* (positive) samples. Their maximal cell-level positive probabilities were only 0.1% and 14.5%, respectively (**Fig. S11**). The false negatives may be because cancer cells were lost in sample processing or not found under the microscope. For POS#7, the patient was diagnosed as negative by the intraoperative frozen section but positive by the postoperative permanent section, so our method had the same finding as the frozen section. The false positive was due to a cell with a 53.4% positive probability, slightly higher than the threshold of 50% (**Fig. S12**). Incorporating more samples, especially those near the MIL-based model’s decision boundary, may help alleviate this problem. Notably, for POS#10 diagnosed as negative by the frozen section but as positive by the permanent section, our method gave out a positive probability of nearly 100%. Despite the frozen section’s high conformity to the permanent section in our study, this is often not the case in medical institutions with limited resources. More clinical data collected from a wider range of medical institutions may be required to evaluate whether our method can surpass the frozen section in low-resource settings and achieve comparable accuracy to that of the permanent section.

The scheme of embedding Raman spectral information acquired by two-color SRS microscopy in this study can be applied to embed multimodal data for more accurate diagnosis, such as extending from two-color to hyperspectral SRS. Solid-state lasers used for SRS microscopy may be replaced with fiber lasers, which offer higher environmental stability, to facilitate clinical translation. Besides, efficiency of sample processing can be improved by utilizing on-chip systems.^53,54^ The imaging throughput can be significantly increased with imaging flow cytometry based on coherent Raman scattering and microfluidic chips, which has realized cell profiling of approximately 100 cells per second.^55^

In summary, we propose a more objective and straightforward approach to intraoperative detection of residual cancer cells on PDAC margins compared to the routine procedure with frozen sections. Our method may find applications in various clinical scenarios, including fine needle aspiration biopsy, ascitic fluid cytology, early cancer detection using bile and pancreatic juice, analysis of circulating tumor cells, and evaluation of minimal residual diseases. Our method’s ability to diagnose with single-cell interpretability, recognize the significant cell subgroup relevant to positive diagnosis, and analyze their abnormal characteristics may make it promising for development into a generalizable tool for cytology.

## ACKNOWLEDGEMENTS

This work was supported by the National Natural Science Foundation of China (No. 62027824).

## AUTHOR INFORMATION

### Contributions

S.Y. and C.H. conceptualized the study. T. F., D. L., and X. C. wrote and edited the paper. All authors reviewed and provided comments on the paper. D. L. provided samples. T. F., X. C., and K. Z. performed experiments and analyzed data. T. F. developed the software. S. Y. and C. H. supervised the study. T. F., D. L., and X. C. contributed equally to this study.

## DATA AVAILABILITY

All data supporting the findings of this study are available within the paper and its supplementary materials. Raw image data and codes are available from the corresponding authors upon reasonable requests.

## METHODS

### Exfoliative Pancreatic Cell Sample Collection

Exfoliative pancreatic cell samples were intraoperatively collected with a medical cytology brush from 41 patients in the Peking University Cancer Hospital. Multiple samples could be collected from the same patient. Collected samples comprised 7 *N*, 28 *T*, 10 *M* (Positive), and 17 *M* (Negative) samples. *N* (normal) samples were collected from seven patients without pancreatic diseases. *T* (tumor) and *M* (PNM) samples were collected from the other 34 patients diagnosed with PDAC. Samples collected from each patient are listed in **Table S2**. Each sample contained ∼1 mL of cell suspension. The inclusion criteria of the 34 patients included: diagnosis as PDAC by permanent section pathology before and after operation; no distant metastasis discovered before and during operation; no multiple primary tumors; no anti-tumor therapy before operation. This study was approved by the Institutional Review Boards of Peking University Cancer Hospital and Beihang University. Written informed consent was obtained from all patients before samples were collected from them and used for our study.

### PDAC Margin Assessment with Frozen Sections and Permanent Sections

During tumor resection, frozen sections were prepared for intraoperative assessment of PNMs. Tissue around the latest PNM was re-resected repeatedly until the resected tissue was found to contain no cancer according to the frozen section. Exfoliative cells used for our study were brushed off from the PNM after the first resection. When the frozen section diagnosis was negative, implying no need for further re-resection, tissue was collected from the eventual PNM and assessed with permanent section pathology after the operation, which is the gold standard of PDAC margin assessment. In the permanent section pathology, a negative PNM was defined according to the 1 mm rule, which demands no cancer cells within 1 mm around the PNM.

In our study, the margin status can be classified into several cases according to the combination of the first frozen section finding and the post-operation permanent section finding (**Table S3**). Most model training labels were specified as the first frozen section finding, but two special cases need to be attended to. When the first frozen section diagnosis was a false negative (which occurred in two cases and meant that the resection ended too early), in the LOOCV, the corresponding *M* sample would be removed from the training set but still used for testing, with the permanent section diagnosis as the ground truth. When re-resection was performed (i.e., the first frozen section diagnosis was positive), the *M* sample corresponded to the first frozen section instead of the permanent section due to different sampling locations. In this case, the first frozen section diagnosis instead of the permanent section diagnosis could serve as the ground truth. For each *M* sample, the frozen and permanent section diagnoses, the maximal COMP#475 value among all cell vectors, and the MICLEAR-predicted patient-level positive probability are listed in **Table S4**.

### Processing of Exfoliative Pancreatic Cell Samples and Two-color SRS Microscopy

Exfoliative pancreatic cell samples were treated with red blood cell lysate for five minutes at 4 °C and then centrifuged at 1200 rpm for 3 minutes. After the supernatant was removed, the cell sediment (∼3 μL) was evenly smeared on a glass slide for SRS microscopy. Our homebuilt SRS microscope employed a dual-output picosecond pulse laser (picoEmeraldTM S, Applied Physics and Electronics) with a repetition rate of 80 MHz and a pulse duration of 2 ps. Its output beams included a pump beam with a tunable wavelength from 700 nm to 960 nm and a Stokes beam with a fixed wavelength at 1031 nm. The Stokes beam was modulated at ∼20 MHz by an electro-optic modulator. Two pulse beams overlapping spatiotemporally were coupled to a 2D scanning galvanometer (GVS012-2D, Thorlabs) and then imported into an inverted microscope (IX73, Olympus). The beams were focused by a 60× water immersion objective (LUMPlanFL N, 1.0 numerical aperture, Olympus) onto the sample and then collected by a 60× water immersion condenser (LUMPlanFL N, 1.0 numerical aperture, Olympus). The Stokes beam was removed by a short-pass filter (ET980SP, Chroma) and then the stimulated Raman loss of the pump beam was detected by a reversely-biased silicon photodiode (S3994-01, Hamamatsu, Japan). The modulated SRS signal was then demodulated by a lock-in amplifier (HF2LI, Zurich Instruments) and read by a data acquisition card (PCIE-6363, National Instruments). A LabVIEW program was written to synchronize galvanometer scanning and SRS signal acquisition. Two-color SRS microscopy was implemented by tuning the pump wavelength to 796.8 nm and 791.8 nm to match the Raman bands at 2850 cm^−1^ and 2930 cm^−1^, corresponding to the C-H stretching region of CH_2_ (rich in lipid) and CH_3_ (rich in protein), respectively. The power at the sample was ∼20 mW and ∼90 mW for the pump and the Stokes beam, respectively. Each SRS image contained 400 × 400 pixels with the field of view ∼100×100 μm. The dwelling time at each pixel was 10 μs.

### Single-cell Segmentation and Quantitative Evaluation

Single-cell segmentation for following machine learning was performed with the Python library *cellpose* 2.2.2. Prediction with the *Cellpose* model requires knowledge of the cell diameter. Smaller cells than this value may be neglected while larger ones may be over-segmented. Because the heterogeneity of cell size can be very high in positive samples, a series of preset values were used to recognize cells of different sizes. The corresponding prediction results were combined to mitigate over-segmentation and loss of cells. After automated single-cell segmentation, few inaccurate results were manually corrected. Single-cell segmentation with watershed and *Stardist* was performed by Fiji and the Python library *stardist 0*.*8*.*3*, respectively. Binary masks for watershed segmentation were created by the Otsu threshold calculated within Fiji.

Performance of segmentation methods was evaluated with three metrics: global Dice coefficient, cell number ratio, and cellwise Dice coefficient. 100 images were randomly chosen for the evaluation, with 50 from *N* and 50 from *T* samples. The ground truth was manual annotation. The global Dice coefficient of each image quantifies the overall overlap degree between the prediction and the ground truth, defined by

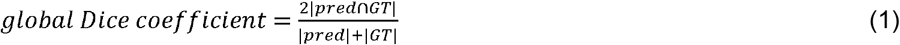

where | · | denotes the overall area of cell regions. The value of the global Dice coefficient is between 0 and 1. Larger values mean better segmentation.

The cell number ratio of each image quantifies the degree of under- or over-segmentation, defined by

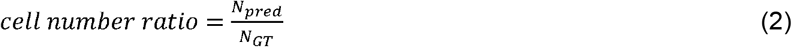

where *N*_*pred*_ and *N*_*GT*_ denote the cell number in the prediction and the ground truth, respectively. The value of the cell number ratio is not less than zero. Values closer to 1 mean better segmentation.

The cellwise Dice coefficient of each image quantifies the mean overlap degree between each segmented cell and the corresponding cell in the ground truth, as defined by

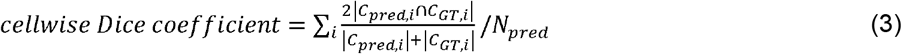

where *C*_*pred*,*i*_ and *C*_*GT*,*i*_ denote the *i -*th cell in the segmentation map and its corresponding cell in the ground truth, respectively. The latter was defined as the cell with the largest intersection over union with the segmented cell. It is an empty set when the largest intersection over union with the segmented cell is 0. The value of the cellwise Dice coefficient is between 0 and 1. Larger values mean better segmentation. This metric is much more sensitive to under- or over-segmentation than the global Dice coefficient.

### Cell Feature Extraction

For each cell, 12 features were manually extracted, including 5 morphological ones (area, perimeter, shape factor, cytoplasm area, and cytoplasm area fraction) and 7 compositional ones (lipid intensity, lipid intensity within cytoplasm, protein intensity, protein intensity within cytoplasm, lipid-protein intensity ratio, LD number, and LD dispersity). The former involves cell shapes and structures. The latter involves intracellular distribution of lipid and protein.

For the pair of two-color SRS images of each field of view, the corresponding molecular image composed of a lipid concentration channel (in green) and a protein concentration channel (in blue) was calculated by

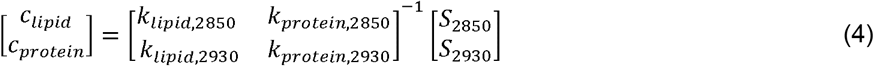

where *k* denotes a composition’s contribution to the SRS signal at a certain Raman shift and was obtained from the Raman spectrum of pure chemical samples; *S* denotes the SRS image pixel intensity at a certain Raman shift; *c* denotes the calculated composition concentration. This formula is derived from the SRS property that SRL is linearly correlated with the concentration of Raman-active vibrational modes. Single-cell molecular images were produced through applying the single-cell segmentation mask to the corresponding molecular image and then placing the segmented cell in the center of a black-background image with 400×400 pixels, which was large enough to contain any cell in the dataset.

Area, centroid, and perimeter were extracted from the segmentation map with the *regionprops* function in the Python library *scikit-image*. Cells on the edge of each field of view were excluded from the feature extraction. Then, the shape factor was calculated by

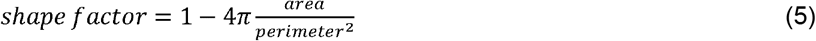

The value of the shape factor is between 0 and 1. The shape factor of a circle is 0. A cell with a slenderer shape has a shape factor closer to 1.

For other features, the feature extraction was performed cell-wise using the segmentation mask. Lipid intensity (mean), protein intensity (mean), and their ratio were calculated with molecular images. LDs were massively detected with the *blob_log* function in the Python library *scikit-image* and then corrected manually. The LD number and LD dispersity (defined as the mean distance from LDs to the cell centroid) were then obtained. Nuclei were detected by applying an Otsu threshold to the lipid channel of each cell because lipid concentration in the nucleus was much lower than that in the cytoplasm. After the nucleus and the cytoplasm were separated, cytoplasm area, cytoplasm area fraction, lipid intensity within cytoplasm (mean), and protein intensity within cytoplasm (mean) were extracted.

### Development of CL-based Cell Embedding Model

The cell embedding model was inspired by a CL framework, *SimCLR*.^38^ CL typically makes a neural network learn how to embed higher-dimensional data, such as spectra and images, into compact lower-dimensional representations by making representations of some data close to each other while others far from each other. *SimCLR* applies certain transformations that do not change the essence of images, such as moderate color distortion. Transformed images of the same image are regarded as similar ones whose low-dimensional representations should be close to each other, while those of different images are regarded as different ones whose low-dimensional representations should be far from each other. This mechanism makes *SimCLR* a self-supervised method. In *SimCLR*, the contrastive loss used to train the embedding model is calculated by

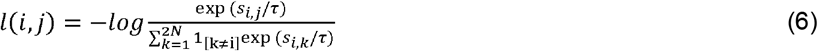

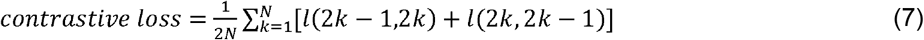

where projections *z*_2*k*-1_ and*z*_2*k*_ came from two transformed images of the *k*-th image in the minibatch containing *N* images (*N*=8 in our case);*s*_*i,j*_ denotes cosine similarity between *z*_*i*_ and *z*_*j*_; *τ* denotes temperature hyperparameter *(τ* =0.07 in our case). A projection was acquired by passing a transformed image through the encoding network *f* and then the smaller projection network *g* .

To obtain the cell embedding model, single-cell molecular images of *N* and *T* samples were divided into a training set and a validation set using stratified sampling with a proportion of 4:1. Images of *M* samples were not used, verifying the model’s capability to embed unseen cells properly. Utilized image transformations included rotation, horizontal flipping, and vertical flipping. Each transformation was performed with a probability of 50%. The cell embedding model was trained with AdamW optimizer for 200 epochs with a NVIDIA GeForce RTX 3070 Laptop GPU. The checkpoint with the smallest contrastive loss on the validation set was chosen. After the model composed of *f* and *g* was trained, *g* was removed for better information preservation.^38^ The cell vector was produced by feeding the single-cell molecular image into the network *f*. The collection of cell vectors of each patient was used for the development of the MIL-based model.

### Development of MIL-based PDAC Margin Assessment Model

The MIL-based model, MI-LR, was developed upon an LR model but trained with an MIL scheme. The LR model’s C parameter was 1.0 and the solver was L-BFGS. The MIL-based model predicted the positive probability of each cell and accordingly gave out the positive probability of each patient. Cells with a positive probability larger than 50% were regarded as significant ones while otherwise as insignificant ones. To initialize MIL-based model training, initial pseudo-labels needed to be assigned to all cells. In *N* and *M* (negative) samples, all cells were assigned negative pseudo-labels. By observing the ISOMAP of cell vectors in *N* and *T* samples, we noticed that there was a significant overlap between them, which might mostly correspond to normal cells that existed in both types of samples. Therefore, we roughly assigned pseudo-labels for cells in *T* and *M* (positive) samples according to their neighbors in the ISOMAP. In the ten nearest neighbors of a cell, if there were fewer than nine belonging to a positive sample (*T* or *M* (positive)), the cell was assigned a negative pseudo-label because its characteristic was considered not unique to cancer cells. Otherwise, the cell was assigned a positive pseudo-label. The LR model was initialized by fitting with these pseudo-labels. Then, cells used for the next training iteration included all cells in negative samples, to which negative pseudo-labels were assigned, and the cell with the largest positive probability in each positive sample, to which positive pseudo-labels were assigned. The LR model was updated after fitting again with these pseudo-labels. The cycle of pseudo-label assignment and model update was repeated until the cell with the largest positive probability in each positive sample stayed unchanged. At this moment, the MIL-based model was considered converged and stopped training. The MI-SVM was trained similarly, but the model was replaced with an SVM classifier with a C parameter of 1.0 and a linear kernel. When changing the parameters of the SVM, we found that in most cases the model failed to converge, so they were not compared with MI-LR. The mi-LR and mi-SVM were different in that in the step of model update, all cells in positive samples were used, with their pseudo-labels coming from the model prediction before the update.

### Statistical Analysis

Dashed lines in violin plots denote medians and two quartiles. Box plots show medians (center lines), two quartiles (top and bottom edges of the box), 2.5 and 97.5 percentiles (top and bottom lines outside the box), and outliers (points). Violin and box plots were drawn with GraphPad Prism 9. Unpaired comparison between cell features in different groups was performed with Mann-Whitney U test, while paired comparison between single-cell segmentation metrics with different segmentation methods was performed with Wilcoxon matched-pairs signed rank test. Two-tailed p-values were calculated in all tests. In violin plots of **Fig. 2d, Fig. 4d**, and **Fig. S4**, outliers were not included because this would compress the violin plot severely, in which outliers were identified with the ROUT method with Q of 0.5%. The removed outliers of **Fig. 2d** were shown in **Fig. S13**. In positive samples, they may be crucial for positive diagnoses but were diluted by excessive cells similar to those in negative samples. The ROC curve and other metrics were calculated by assembling 27 PDAC margin assessment results in the LOOCV, in which predicted positive probabilities, patient-level prediction results, and ground truths were used for calculation.

