## Supplementary Materials for "MICLEAR: Intelligent Molecular Cytology for Intraoperative Margin Assessment of Pancreatic Ductal Adenocarcinoma"

^1^Key Laboratory of Biomechanics and Mechanobiology (Beihang University), Ministry of Education; Key Laboratory of Innovation and Transformation of Advanced Medical Devices, Ministry of Industry and Information Technology; National Medical Innovation Platform for Industry-Education Integration in Advanced Medical Devices (Interdiscipline of Medicine and Engineering); School of Biological Science and Medical Engineering, Beihang University, Beijing, 100191, China.

^2^Key laboratory of Carcinogenesis and Translational Research (Ministry of Education/Beijing), Department of Hepato-Pancreato-Biliary Surgery/Sarcoma Center, Peking University Cancer Hospital & Institute, Beijing, 100142, China.

^3^School of Engineering Medicine, Beihang University, Beijing, 100191, China.

^#^These authors contributed equally.

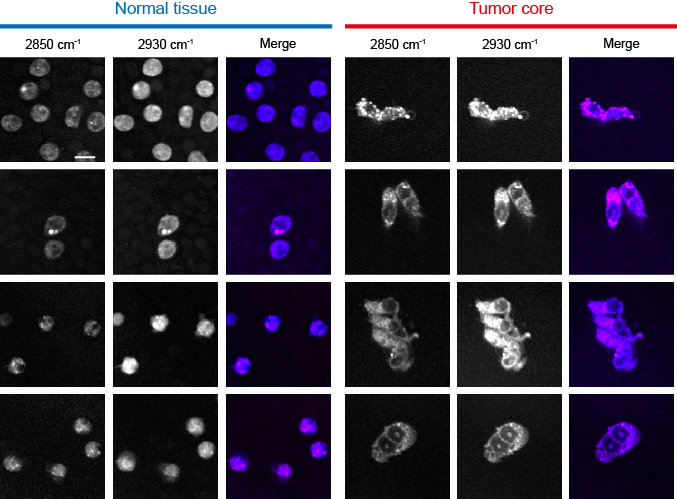

**Figure S1 | Raw SRS images in *N* and *T* samples.** Raw SRS images were acquired at 2850 cm^-1^ (magenta channel in merged images) and 2930 cm^-1^ (blue channel in merged images). Scale bars, 10 μm.

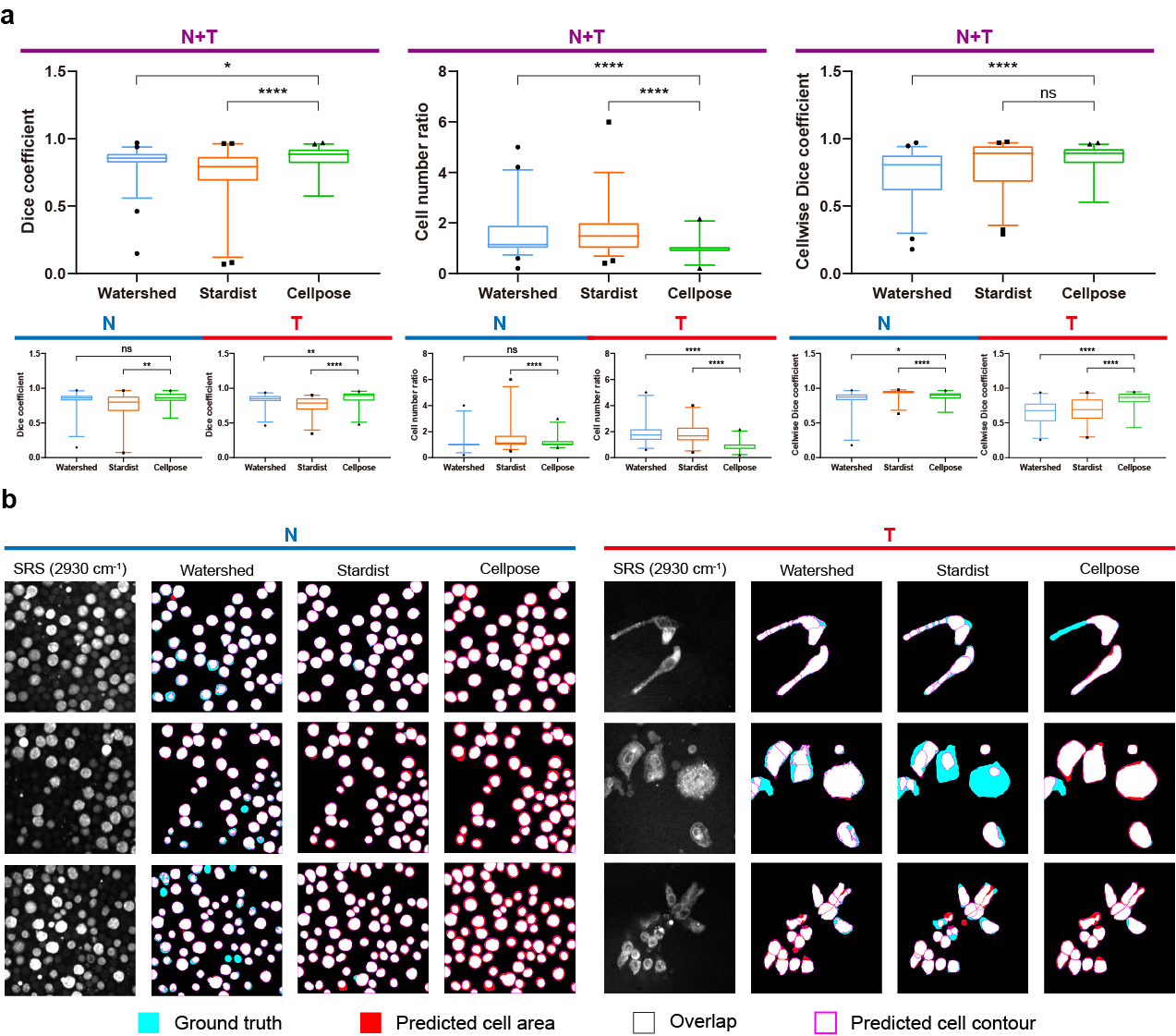

**Figure S2 | Single-cell segmentation with *Cellpose*. a**, Performance of segmentation with three methods on randomly chosen 50 *N* and 50 *T* sample images. SRS images at 2930 cm^-1^ were used for segmentation. The difference between the performance of *Cellpose* and the other two methods was more obvious when segmenting cells in *T* sample images. **b**, Representative segmentation results with three methods. Watershed frequently resulted in over-segmentation. *Stardist* performed poorly on nonconvex cells. *Cellpose* performed well on segmentation of both convex and nonconvex cells. ns: no significant difference, *: p<0.05, **: p<0.01, ****: p<0.0001; tested by Mann-Whitney U test.

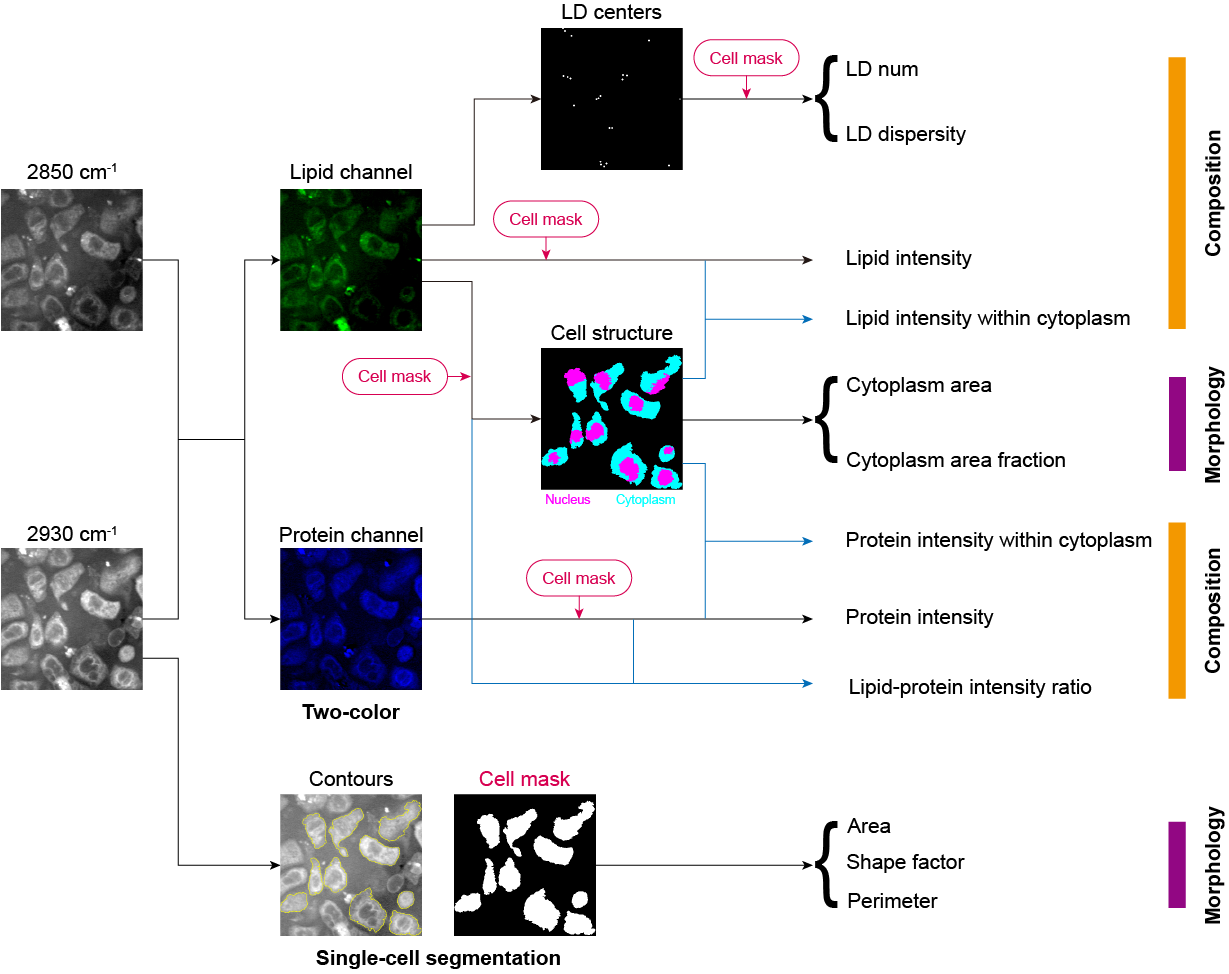

**Figure S3 | Workflow of cell feature extraction.** Cell masks were obtained with 2930 cm^-1^ images for the following feature extraction. Three of morphological features (area, shape factor, and perimeter) were directly extracted from cell masks. Distribution of lipid and protein in each cell was calculated with SRS images at 2850 cm^-1^ and 2930 cm^-1^, and then three of compositional features (lipid intensity, protein intensity, and lipid-protein intensity ratio) were extracted. LDs were detected with the lipid channel and then two of compositional features (LD num and LD dispersity) were extracted. Finally, nucleus and cytoplasm were separated with the lipid channel and then two of morphological features (cytoplasm area and cytoplasm area faction) and two of compositional features (lipid intensity within cytoplasm and protein intensity within cytoplasm) were extracted.

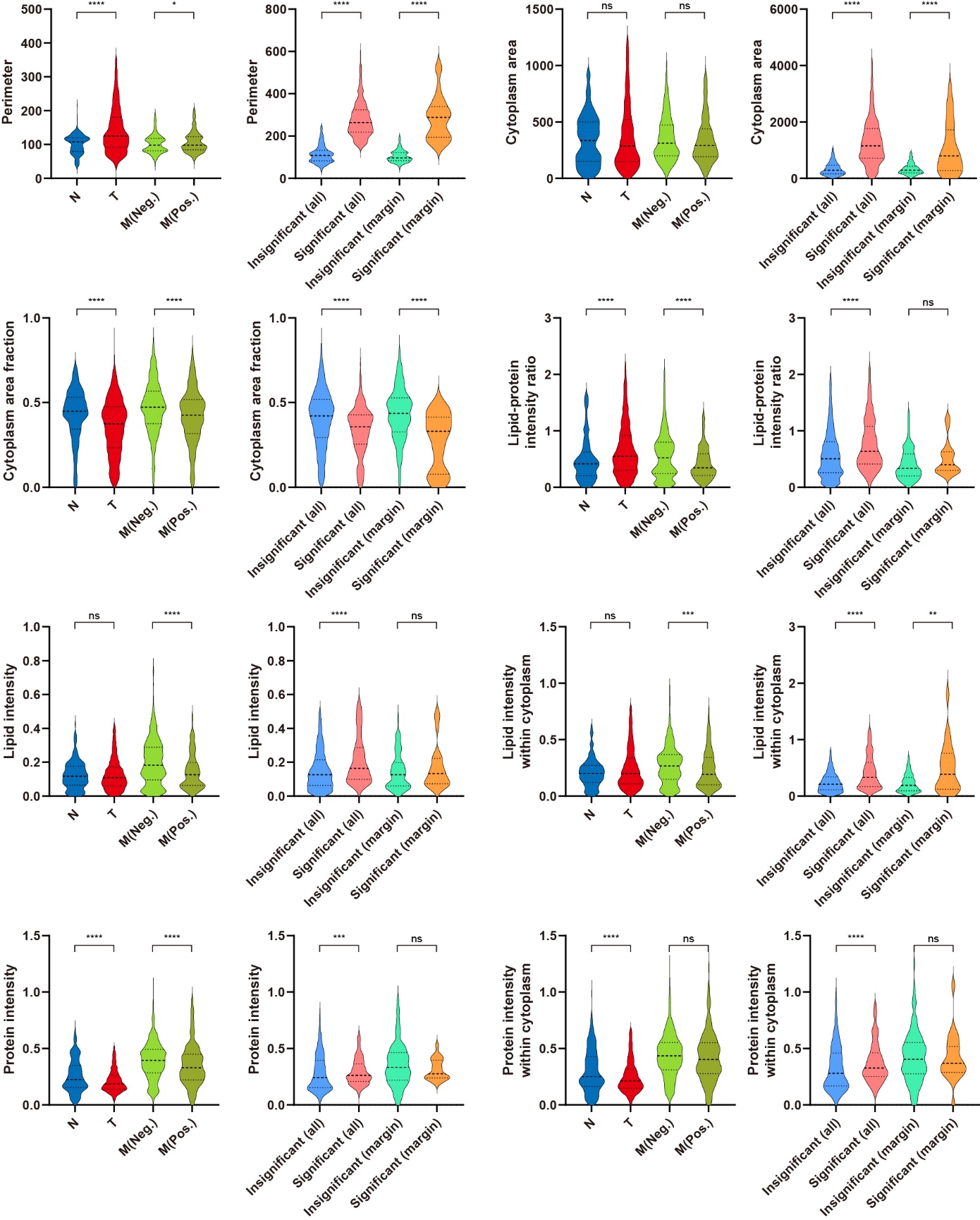

**Figure S4 | Comparison of cell features before and after significant cell recognition by the MIL-based model.** ns: no significant difference, *: p<0.05; **: p<0.01, ***: p<0.001, ****: p<0.0001; tested by Mann-Whitney U test.

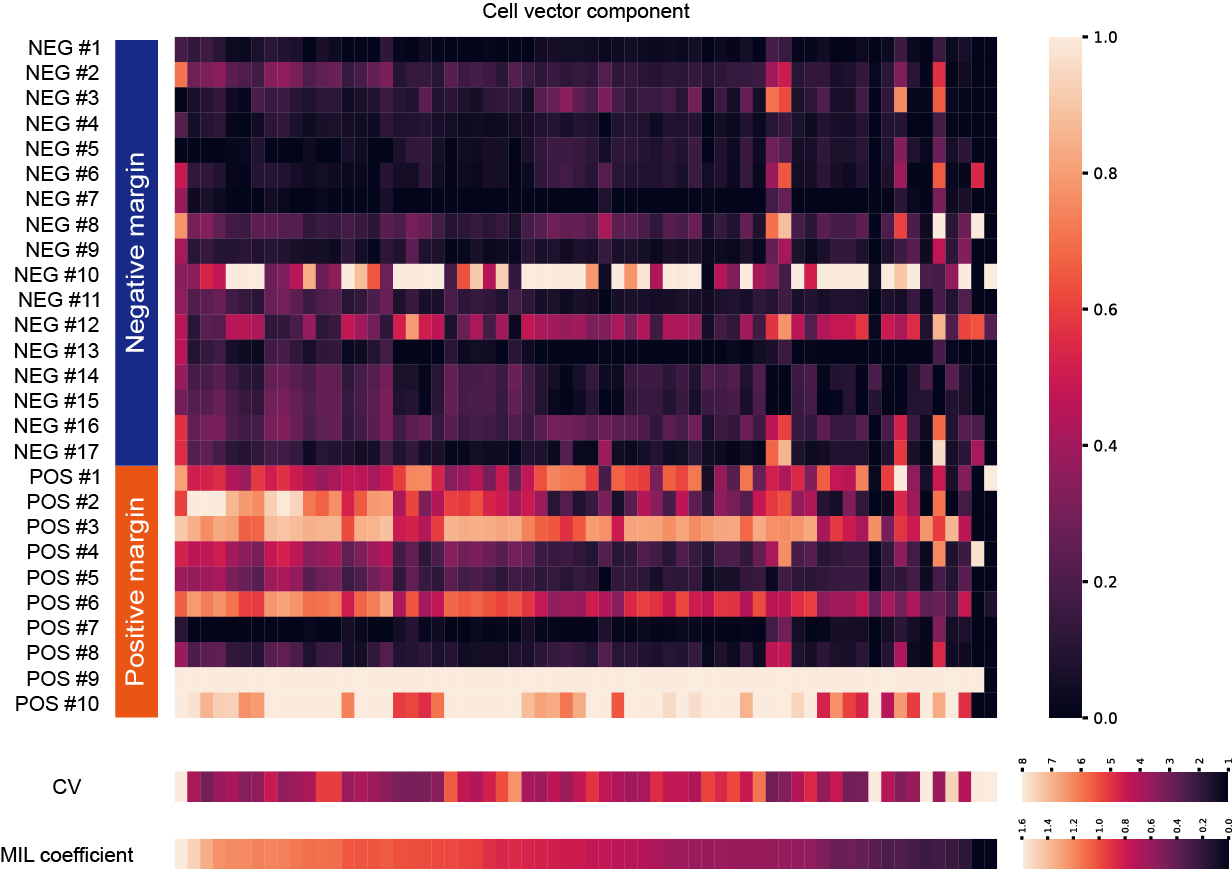

**Figure S5 | Heatmap of cell vector component maximums in *M* samples.** Each column denotes a cell vector component. Each row denotes an *M* sample. Each grid denotes the maximum of a certain cell vector component among all cells in a sample. Maximums of each cell vector component were normalized between 0 and 1. Only cell vector components with CV larger than three are presented here.

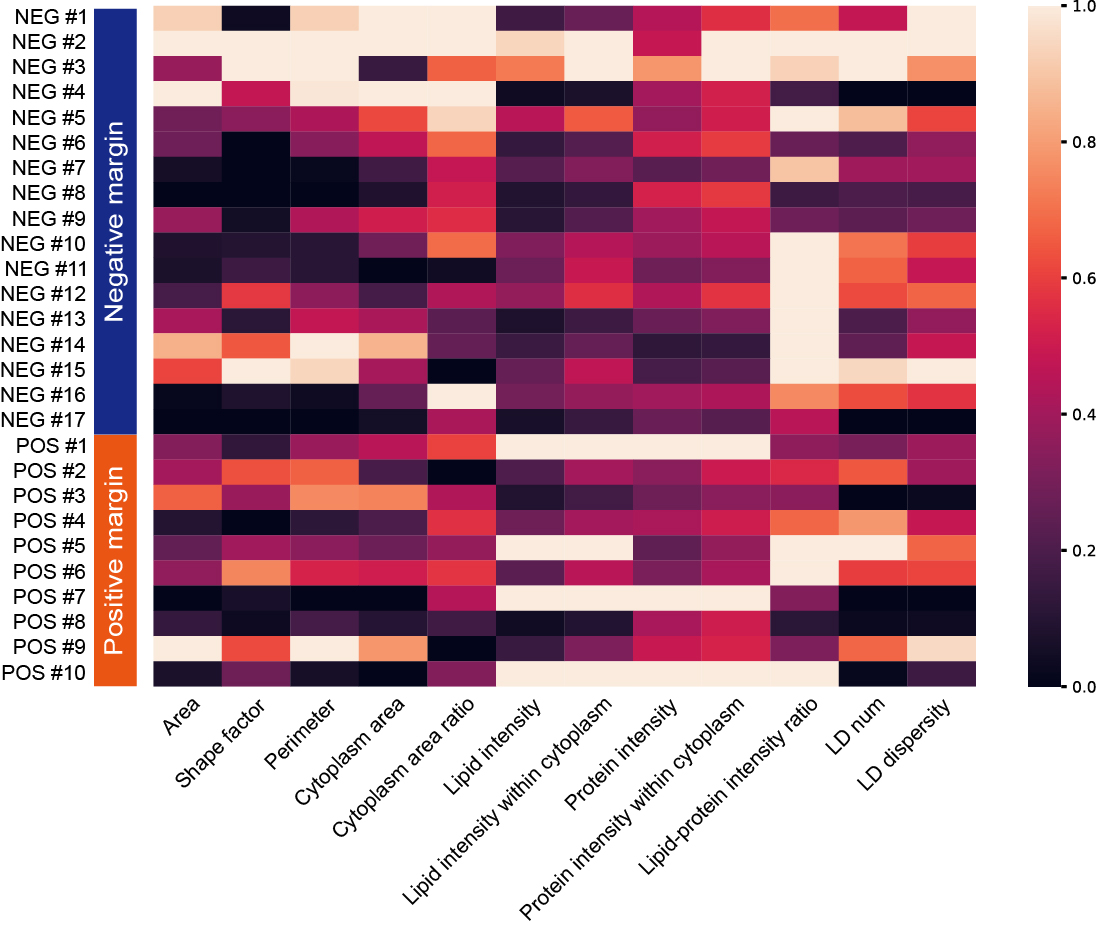

**Figure S6 | Heatmap of mean feature values in *M* samples.** Mean values of each extracted cell feature were normalized between 0 and 1.

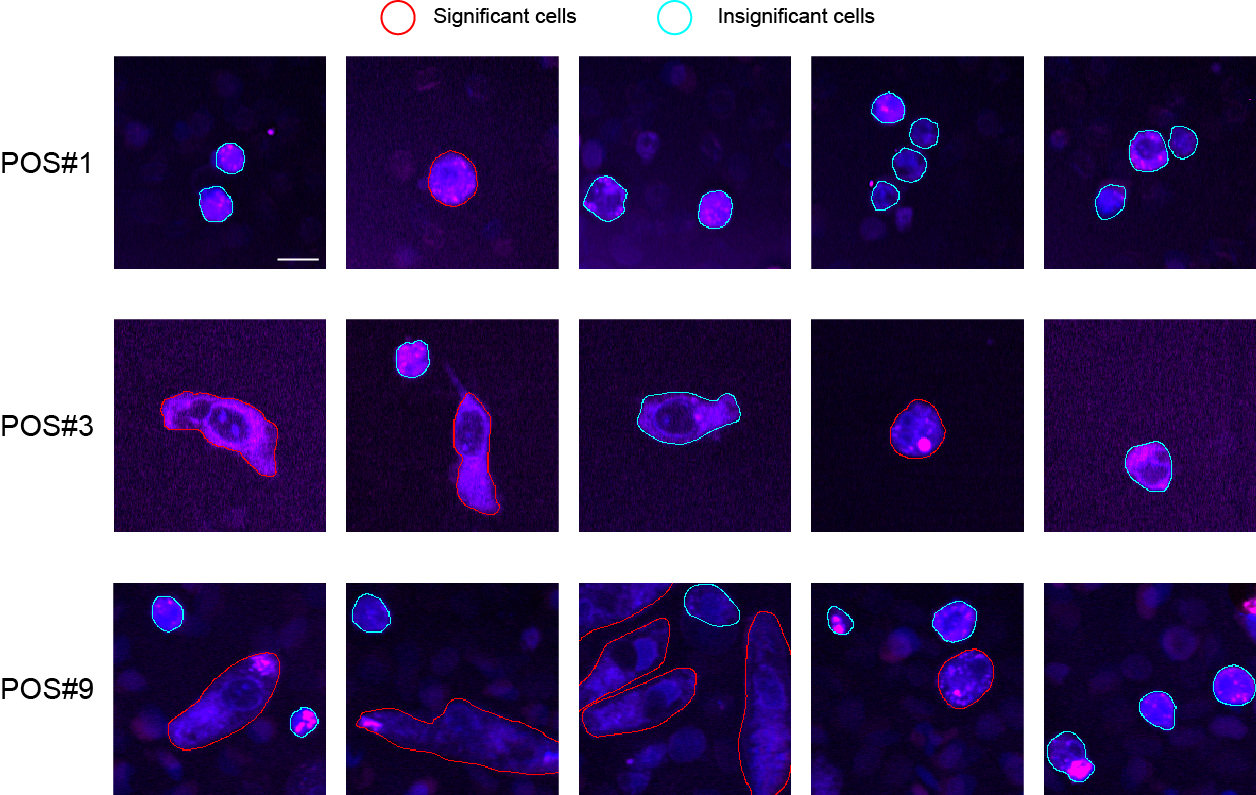

**Figure S7 | Recognized significant cells in *M* (positive) samples.** Cells with positive probability larger than 0.5 were termed as significant cells and marked with red contours. The 2850 cm^-1^ image was in the magenta channel and the 2930 cm^-1^ image was in the blue channel. Scale bars, 10 μm.

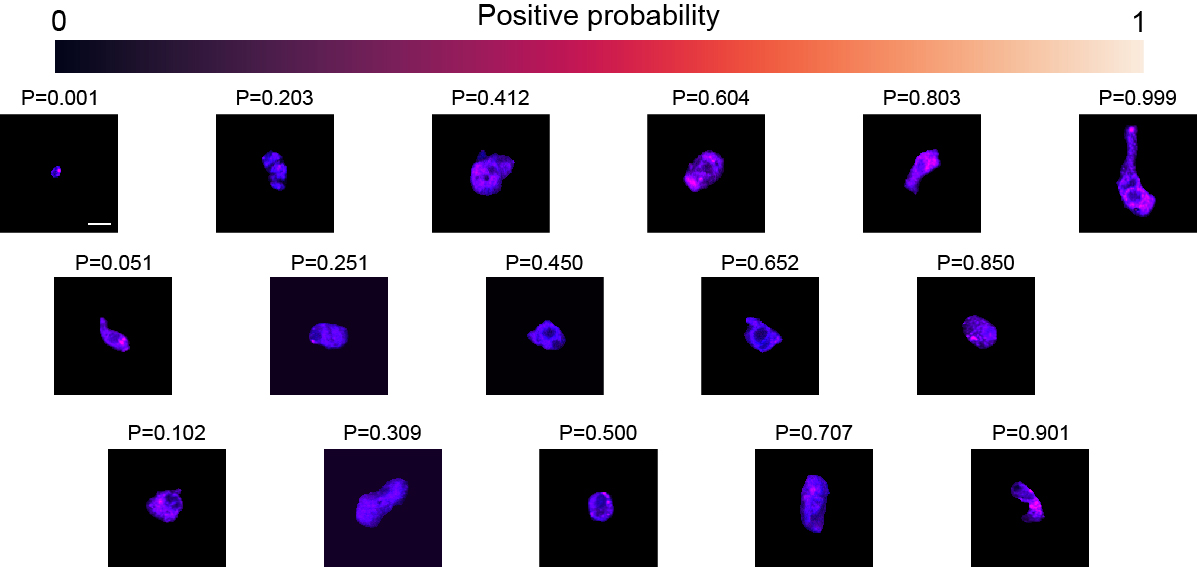

**Figure S8 | Examples of cells and corresponding positive probabilities.** Here, probabilities were outputted by the MI-LR model whose coefficients were the average of all models obtained in LOOCV folds. The 2850 cm^-1^ image was in the magenta channel and the 2930 cm^-1^ image was in the blue channel. Scale bars, 10 μm.

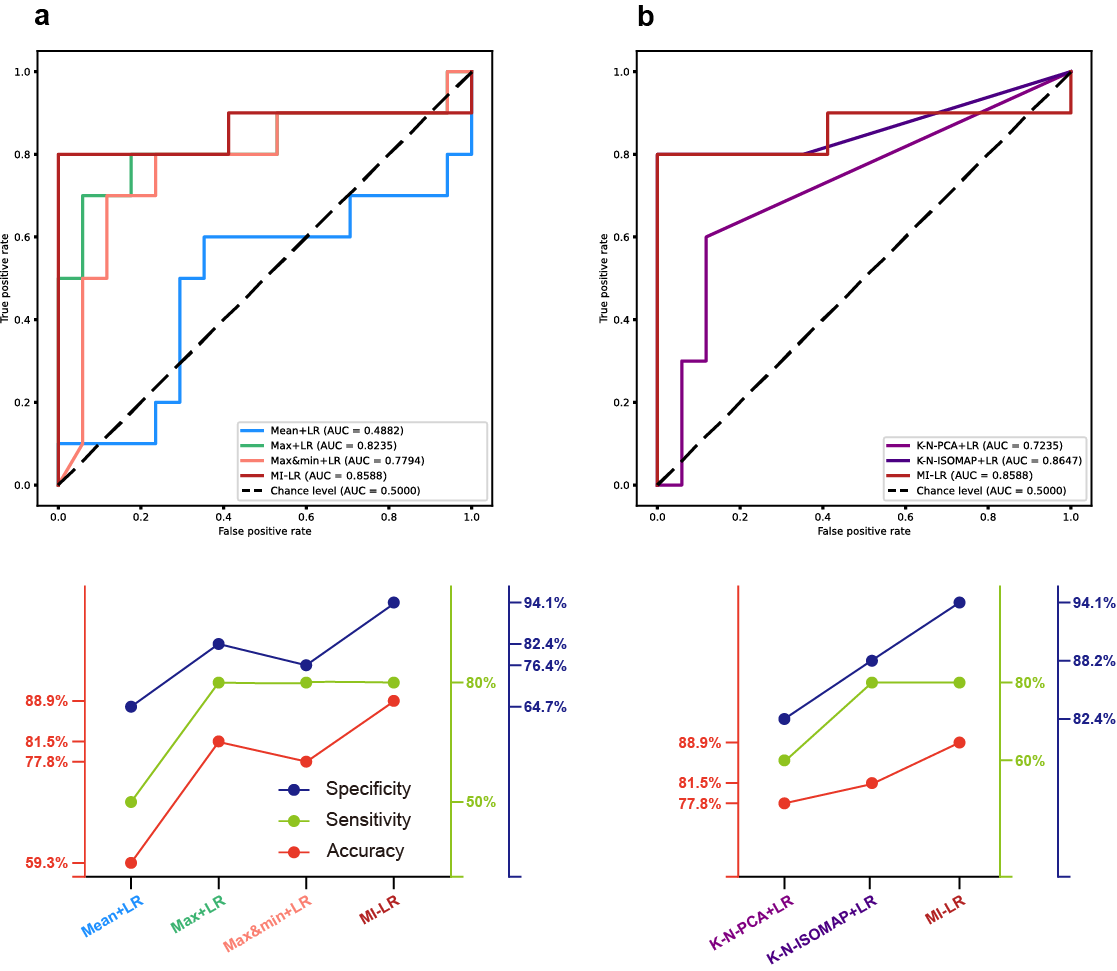

**Figure S9 | Performance of the MI-LR model and other machine learning algorithms using LR. a**, Comparison with methods using sample-level statistics for margin assessment. **b**, Comparison with methods using clustering for margin assessment. All methods used the cell vectors produced by the cell embedding model.

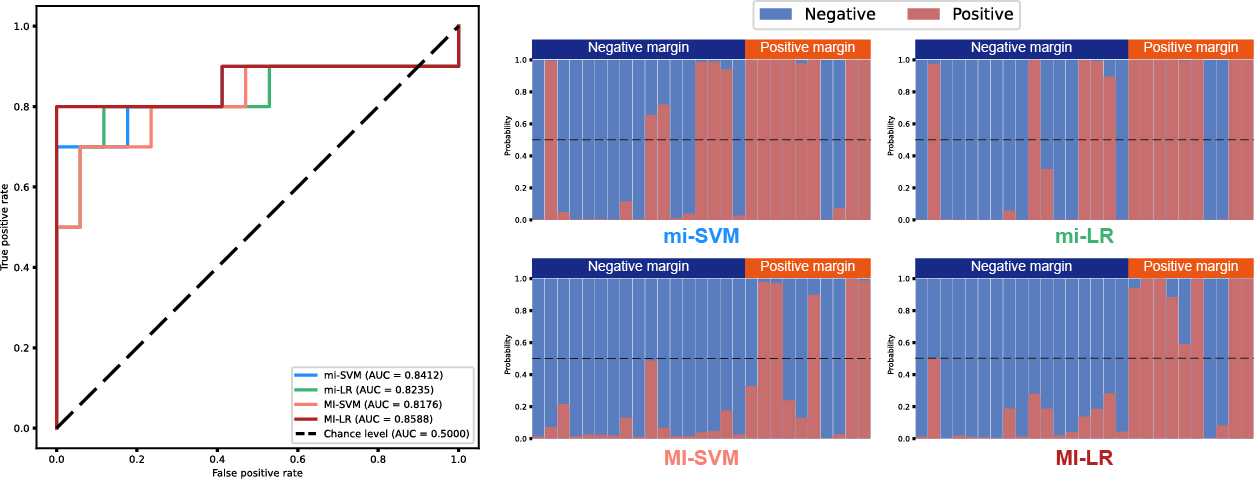

**Figure S10 | Performance of the MI-LR model and other MIL models.** All methods used the cell vectors produced by the cell embedding model.

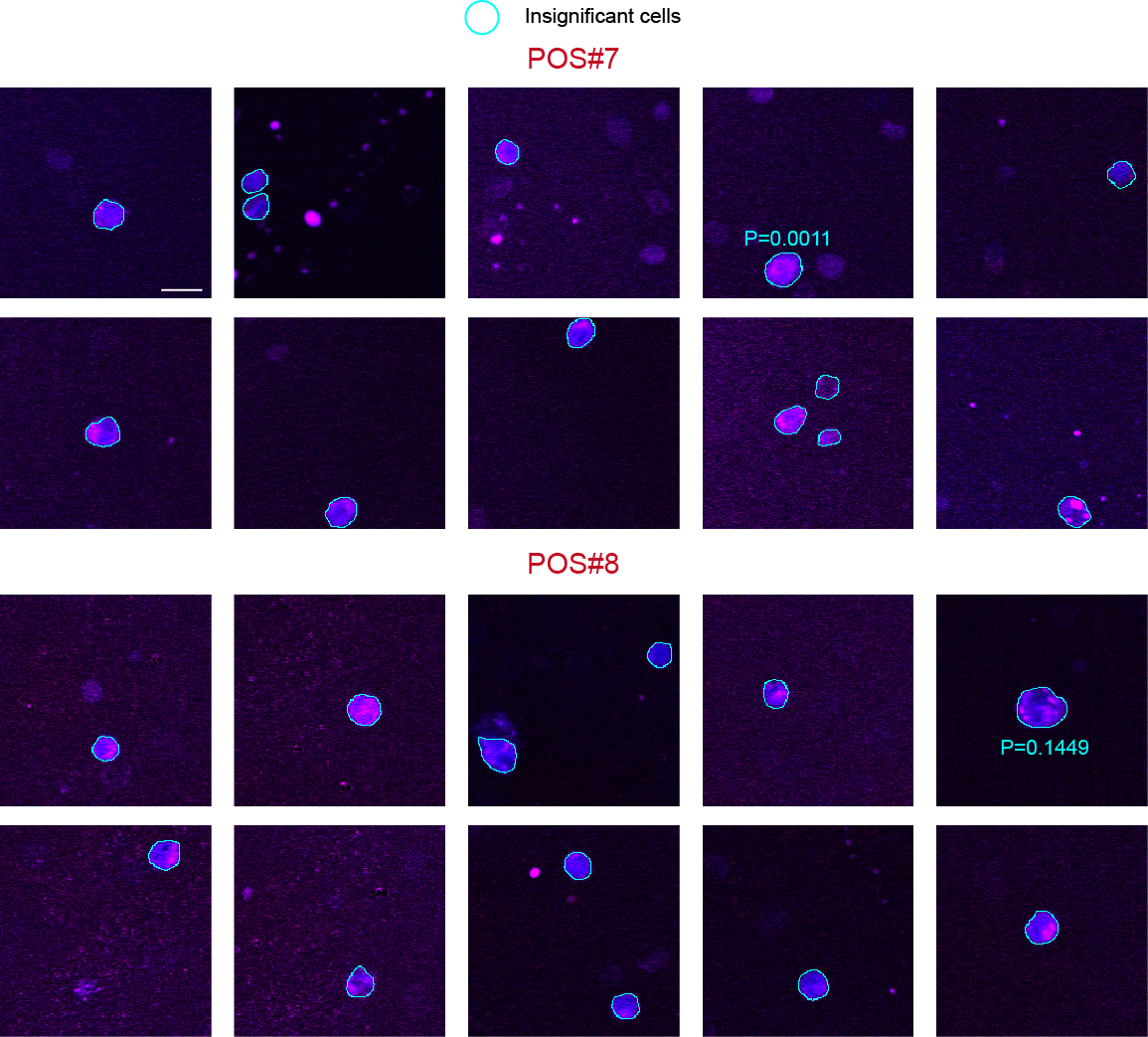

**Figure S11 | False negatives in PDAC margin assessment.** The cell with the largest positive probability in the sample was marked with its probability value. The 2850 cm^-1^ image was in the magenta channel and the 2930 cm^-1^ image was in the blue channel. Scale bars, 10 μm.

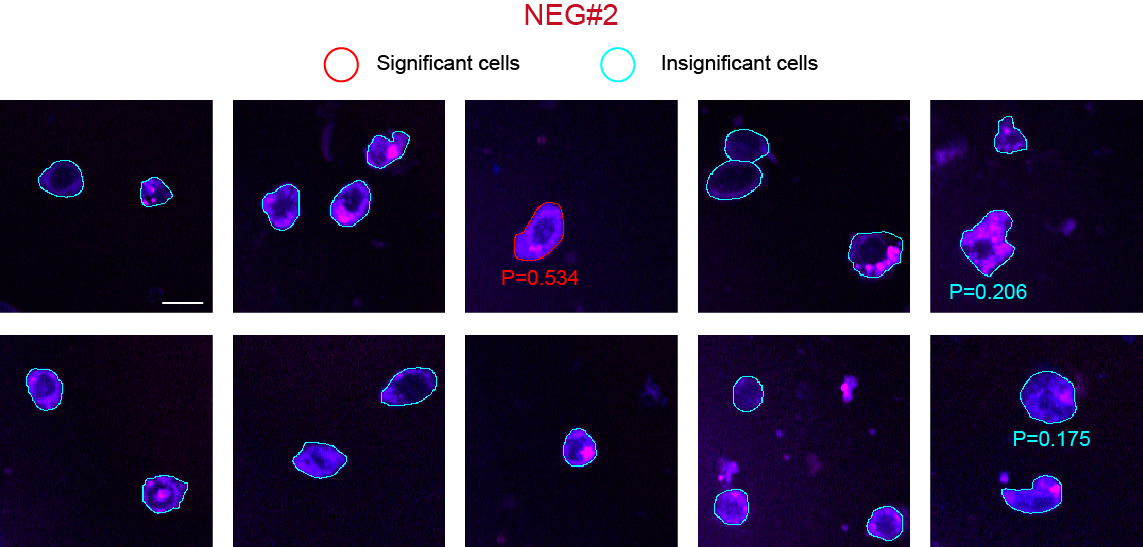

**Figure S12 | False positive in PDAC margin assessment.** Cells with positive probability larger than 0.1 in the sample were marked with its probability value. The 2850 cm^-1^ image was in the magenta channel and the 2930 cm^-1^ image was in the blue channel. Scale bars, 10 μm.

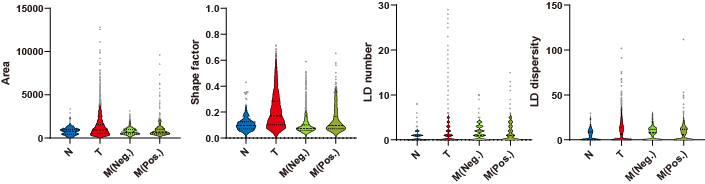

**Figure S13 | Feature distribution in different sample groups without removing outliers.** Gray points denote outliers removed in **Fig. 2d**.

**Table S1 | MIL model coefficients, CVs, and SCCs of cell vector components, sorted by the MIL coefficient in an ascending order.**

| Cell vector component | MIL model coefficient | CV | SCC with area | SCC with shape factor | SCC with LD number | SCC with LD dispersity |
| --- | --- | --- | --- | --- | --- | --- |
| #0 | -7.15212 | 0.02453 | 0.62482 | 0.34189 | 0.17479 | 0.5476 |
| #1 | -6.86464 | 0.09925 | 0.93213 | 0.44042 | 0.38216 | 0.82766 |
| #2 | -5.35033 | 0.82596 | 0.1898 | 0.00235 | 0.10028 | 0.16957 |
| #3 | -4.47606 | 0.02829 | 0.3775 | 0.3638 | 0.06407 | 0.33654 |
| #4 | -4.1771 | 0.01149 | -0.33293 | -0.23229 | -0.1092 | -0.30083 |
| #5 | -4.42516 | 0.01806 | 0.5944 | 0.33484 | 0.17171 | 0.52866 |
| #6 | -3.9009 | 0.0051 | 0.36081 | 0.02538 | 0.21215 | 0.31158 |
| #7 | -3.99891 | 0.02 | 0.33829 | 0.13338 | 0.10752 | 0.30027 |
| #8 | -3.871 | 0.01307 | -0.08768 | -0.14781 | -0.06309 | -0.07923 |
| #9 | -3.86329 | 0.00614 | -0.06713 | -0.26601 | 0.02138 | -0.07243 |
| #10 | -3.27779 | 0.00925 | 0.14152 | -0.07768 | 0.08061 | 0.1216 |
| #11 | -3.59706 | 0.05663 | -0.14871 | -0.26607 | -0.04549 | -0.13827 |
| #12 | -3.36503 | 0.14635 | -0.02337 | -0.23583 | 0.05899 | -0.02373 |
| #13 | -3.25432 | 0.04749 | -0.00322 | -0.21724 | 0.05889 | -0.00886 |
| #14 | -3.1478 | 0.00842 | 0.05915 | -0.12158 | 0.03518 | 0.04776 |
| #15 | -3.25515 | 1.40435 | 0.20182 | -0.14562 | 0.15256 | 0.16799 |
| #16 | -3.14653 | 0.05465 | 0.15196 | -0.1236 | 0.11418 | 0.1253 |
| #17 | -2.81817 | 0.04183 | -0.14217 | -0.22544 | -0.02295 | -0.12828 |
| #18 | -2.71533 | 0.00946 | 0.07589 | -0.11901 | 0.09442 | 0.06285 |
| #19 | -2.77206 | 0.02358 | -0.00478 | -0.13878 | 0.00929 | -0.00299 |
| #20 | -2.46847 | 0.62608 | -0.11994 | -0.28806 | 0.13836 | -0.12841 |
| #21 | -2.15803 | 0.02656 | 0.13067 | 0.04024 | 0.09609 | 0.11968 |
| #22 | -2.49151 | 0.01702 | 0.05103 | -0.13431 | 0.02764 | 0.03402 |
| #23 | -2.19858 | 0.00699 | -0.09096 | -0.19046 | -0.02012 | -0.08948 |
| #24 | -2.12925 | 1.31269 | 0.20408 | -0.16346 | 0.20265 | 0.17144 |
| #25 | -2.00234 | 1.40305 | -0.66482 | -0.31087 | -0.30029 | -0.60402 |
| #26 | -2.06048 | 1.23831 | -0.35074 | -0.48926 | -0.02277 | -0.3224 |
| #27 | -1.92712 | 0.02824 | -0.38816 | -0.37367 | -0.11707 | -0.35065 |
| #28 | -1.95217 | 1.09147 | -0.69543 | -0.5832 | -0.17704 | -0.6319 |
| #29 | -2.00374 | 0.03485 | 0.34654 | 0.07578 | 0.15508 | 0.31318 |
| #30 | -1.74497 | 1.40749 | -0.78396 | -0.38576 | -0.35083 | -0.70888 |
| #31 | -1.76599 | 0.02314 | 0.17161 | -0.04872 | 0.06884 | 0.1429 |
| #32 | -1.44563 | 0.04767 | 0.54949 | 0.10017 | 0.29912 | 0.48389 |
| #33 | -1.67765 | 0.01523 | 0.10267 | -0.08311 | 0.02878 | 0.07974 |
| #34 | -1.45314 | 1.70779 | -0.80222 | -0.50369 | -0.29976 | -0.71861 |
| #35 | -1.50269 | 1.05452 | -0.54178 | -0.55905 | -0.10579 | -0.49464 |
| #36 | -1.52988 | 1.32459 | -0.01982 | -0.15717 | 0.02859 | -0.0345 |
| #37 | -1.48527 | 1.00126 | -0.53943 | -0.55647 | -0.11208 | -0.49259 |
| #38 | -1.39009 | 1.24058 | -0.37226 | -0.49295 | -0.00598 | -0.33941 |
| #39 | -1.407 | 2.13978 | 0.04532 | -0.20105 | 0.21608 | 0.03198 |
| #40 | -1.32501 | 1.92558 | -0.66115 | -0.33929 | -0.23489 | -0.59859 |
| #41 | -1.40263 | 1.10914 | -0.66354 | -0.56361 | -0.16423 | -0.59918 |
| #42 | -1.24523 | 1.85698 | -0.1751 | 0.12421 | -0.11317 | -0.15079 |
| #43 | -1.34635 | 0.98129 | -0.42505 | -0.53086 | -0.05006 | -0.39377 |
| #44 | -0.75317 | 0.00451 | 0.61915 | 0.1319 | 0.37803 | 0.54546 |
| #45 | -1.31732 | 1.07847 | -0.63885 | -0.582 | -0.16123 | -0.58088 |
| #46 | -1.26318 | 0.00503 | 0.43736 | 0.10577 | 0.18881 | 0.37878 |
| #47 | -1.2297 | 1.54044 | -0.70544 | -0.43717 | -0.22773 | -0.63361 |
| #48 | -1.25598 | 1.59875 | 0.31074 | -0.03278 | 0.18298 | 0.26881 |
| #49 | -1.28173 | 0.00839 | 0.06754 | -0.09557 | 0.01834 | 0.04709 |
| #50 | -1.14819 | 1.86215 | -0.70085 | -0.3782 | -0.25695 | -0.62471 |
| #51 | -1.1512 | 1.88327 | -0.61602 | -0.31516 | -0.2219 | -0.55446 |
| #52 | -1.16242 | 1.86491 | -0.77363 | -0.4293 | -0.27898 | -0.6907 |
| #53 | -1.1678 | 1.65002 | -0.84862 | -0.41732 | -0.34022 | -0.75843 |
| #54 | -1.12616 | 1.88782 | -0.72962 | -0.38018 | -0.26185 | -0.65448 |
| #55 | -1.11561 | 1.67167 | -0.77257 | -0.46808 | -0.26979 | -0.69814 |
| #56 | -1.13249 | 1.73235 | -0.78058 | -0.41049 | -0.26887 | -0.69922 |
| #57 | -1.17338 | 1.20701 | -0.74202 | -0.58243 | -0.22284 | -0.6736 |
| #58 | -1.10586 | 1.67783 | -0.81526 | -0.51554 | -0.2744 | -0.73601 |
| #59 | -1.10788 | 1.44096 | -0.73196 | -0.54085 | -0.21862 | -0.66833 |
| #60 | -1.11869 | 1.77684 | -0.7561 | -0.47755 | -0.24525 | -0.68106 |
| #61 | -1.07606 | 1.88021 | -0.56337 | -0.20591 | -0.18398 | -0.49628 |
| #62 | -1.16563 | 1.31564 | 0.12436 | -0.11567 | 0.10735 | 0.09652 |
| #63 | -1.00682 | 0.8496 | -0.81122 | -0.29077 | -0.41111 | -0.73005 |
| #64 | -1.11181 | 1.19797 | -0.68631 | -0.57943 | -0.18909 | -0.62389 |
| #65 | -1.0003 | 1.30247 | -0.18317 | 0.11677 | -0.11925 | -0.16066 |
| #66 | -1.04494 | 1.74615 | -0.80649 | -0.4464 | -0.29854 | -0.72831 |
| #67 | -1.01939 | 1.6803 | -0.73112 | -0.50932 | -0.22578 | -0.66199 |
| #68 | -1.02436 | 1.73805 | -0.66873 | -0.27722 | -0.2676 | -0.6031 |
| #69 | -0.99749 | 1.57316 | -0.67847 | -0.46414 | -0.20783 | -0.61388 |
| #70 | -1.04649 | 1.01291 | -0.47949 | -0.528 | -0.07966 | -0.43676 |
| #71 | -1.008 | 1.76973 | -0.82181 | -0.37081 | -0.3286 | -0.73757 |
| #72 | -0.98707 | 1.70794 | -0.85516 | -0.51571 | -0.28759 | -0.76956 |
| #73 | -0.98046 | 1.81954 | -0.81754 | -0.4376 | -0.29026 | -0.73193 |
| #74 | -0.98336 | 1.79493 | -0.79858 | -0.37259 | -0.30089 | -0.71702 |
| #75 | -1.06515 | 0.01975 | 0.19651 | -0.03306 | 0.08717 | 0.16946 |
| #76 | -0.95799 | 1.52196 | -0.64487 | -0.4126 | -0.20944 | -0.58419 |
| #77 | -0.97204 | 1.59416 | -0.89928 | -0.51645 | -0.33859 | -0.80503 |
| #78 | -0.97094 | 1.23224 | -0.44803 | -0.50104 | -0.0573 | -0.40484 |
| #79 | -0.96048 | 1.11072 | -0.502 | -0.56746 | -0.07309 | -0.46284 |
| #80 | -0.95326 | 1.72745 | -0.84983 | -0.52145 | -0.29277 | -0.76459 |
| #81 | -0.99704 | 0.95466 | -0.62807 | -0.58096 | -0.15031 | -0.57346 |
| #82 | -0.94098 | 1.7446 | -0.80198 | -0.49695 | -0.25905 | -0.72302 |
| #83 | -0.88203 | 2.11628 | 0.26151 | -0.12374 | 0.24705 | 0.23367 |
| #84 | -0.75342 | 0.00521 | 0.69498 | 0.3284 | 0.3644 | 0.62503 |
| #85 | -0.91403 | 1.66754 | -0.83631 | -0.52569 | -0.27744 | -0.75415 |
| #86 | -0.95616 | 0.89038 | -0.4968 | -0.55505 | -0.08156 | -0.45785 |
| #87 | -0.91993 | 1.06618 | -0.53601 | -0.55923 | -0.09298 | -0.49114 |
| #88 | -0.9394 | 1.20591 | -0.72902 | -0.57584 | -0.21905 | -0.6602 |
| #89 | -0.77569 | 0.39017 | 0.34876 | 0.38669 | 0.07321 | 0.30845 |
| #90 | -0.89123 | 1.42549 | -0.8641 | -0.5453 | -0.29827 | -0.77672 |
| #91 | -1.08602 | 0.02847 | -0.55214 | -0.39381 | -0.23233 | -0.50009 |
| #92 | -0.8915 | 1.72315 | -0.66228 | -0.30625 | -0.21536 | -0.5859 |
| #93 | -0.90462 | 1.14015 | -0.61527 | -0.54669 | -0.11776 | -0.55811 |
| #94 | -0.92448 | 0.97214 | -0.6021 | -0.55772 | -0.14565 | -0.54579 |
| #95 | -0.90278 | 1.15542 | -0.64607 | -0.56609 | -0.16932 | -0.58626 |
| #96 | -0.84786 | 1.56062 | -0.57675 | -0.27308 | -0.17337 | -0.51142 |
| #97 | -0.8705 | 0.9612 | -0.53381 | -0.562 | -0.09829 | -0.49017 |
| #98 | -0.84183 | 1.67928 | -0.77538 | -0.43665 | -0.27334 | -0.69727 |
| #99 | -0.80805 | 0.7267 | 0.03465 | -0.30568 | 0.17742 | 0.01565 |
| #100 | -0.68946 | 0.00596 | 0.63337 | 0.453 | 0.14555 | 0.55894 |
| #101 | -0.7798 | 1.42662 | -0.85895 | -0.47139 | -0.30936 | -0.77143 |
| #102 | -0.78244 | 1.20312 | -0.58248 | -0.53505 | -0.11775 | -0.5252 |
| #103 | -0.77781 | 0.00574 | 0.20098 | -0.03245 | 0.09727 | 0.16664 |
| #104 | -0.7499 | 1.5736 | -0.85273 | -0.45799 | -0.34473 | -0.76069 |
| #105 | -0.78361 | 1.17529 | -0.64559 | -0.5624 | -0.17435 | -0.58445 |
| #106 | -0.6984 | 0.00863 | 0.42533 | 0.05999 | 0.24066 | 0.36492 |
| #107 | -0.79954 | 0.02221 | -0.76602 | -0.30365 | -0.44268 | -0.68421 |
| #108 | -0.70422 | 1.11009 | -0.56592 | -0.56996 | -0.10881 | -0.52004 |
| #109 | -0.63181 | 2.1922 | -0.12755 | -0.34343 | 0.08956 | -0.11679 |
| #110 | -0.66246 | 1.63283 | -0.44928 | -0.28685 | -0.09039 | -0.40133 |
| #111 | -0.67075 | 1.06665 | -0.49287 | -0.5162 | -0.02803 | -0.44756 |
| #112 | -0.63003 | 0.01569 | -0.02102 | -0.03246 | -0.05694 | -0.02679 |
| #113 | -0.64471 | 1.04198 | -0.57941 | -0.57231 | -0.12501 | -0.53061 |
| #114 | -0.63443 | 1.33211 | 0.34865 | -0.05299 | 0.23632 | 0.29954 |
| #115 | -0.63983 | 1.43455 | 0.13033 | -0.17042 | 0.1389 | 0.10686 |
| #116 | -0.54449 | 2.37058 | -0.07281 | -0.2941 | 0.08676 | -0.0637 |
| #117 | -0.65102 | 1.033 | -0.63508 | -0.56717 | -0.1632 | -0.57798 |
| #118 | -0.61709 | 0.94031 | -0.4117 | -0.52782 | -0.04136 | -0.38216 |
| #119 | -0.6369 | 0.90254 | -0.43595 | -0.54209 | -0.03848 | -0.40539 |
| #120 | -0.61618 | 1.3689 | -0.75676 | -0.47267 | -0.21588 | -0.68077 |
| #121 | -0.64434 | 0.91371 | -0.54365 | -0.56323 | -0.10195 | -0.49832 |
| #122 | -0.59821 | 1.54694 | 0.22348 | -0.12991 | 0.19228 | 0.1878 |
| #123 | -0.62241 | 1.44281 | 0.25821 | -0.09186 | 0.19783 | 0.22224 |
| #124 | -0.5497 | 1.77337 | -0.28626 | -0.38183 | 0.00168 | -0.25288 |
| #125 | -0.60381 | 0.97719 | -0.47532 | -0.54721 | -0.07196 | -0.4386 |
| #126 | -0.53962 | 0.00322 | 0.48059 | 0.09605 | 0.21716 | 0.42285 |
| #127 | -0.59335 | 1.50205 | 0.13343 | -0.13092 | 0.12085 | 0.11201 |
| #128 | -0.62581 | 1.01438 | -0.70856 | -0.56028 | -0.1979 | -0.64161 |
| #129 | -0.52997 | 1.31327 | 0.25796 | -0.12257 | 0.21701 | 0.22016 |
| #130 | -0.56814 | 1.02231 | -0.67778 | -0.55948 | -0.18446 | -0.61271 |
| #131 | -0.46547 | 2.3328 | 0.20483 | -0.1537 | 0.23398 | 0.18249 |
| #132 | -0.53645 | 0.96695 | -0.35352 | -0.50887 | -0.009 | -0.33095 |
| #133 | -0.53967 | 1.49059 | 0.17858 | -0.15776 | 0.16541 | 0.14947 |
| #134 | -0.52464 | 0.92738 | -0.25299 | -0.47298 | 0.04961 | -0.24328 |
| #135 | -0.48891 | 2.13567 | 0.13448 | -0.1791 | 0.18608 | 0.1212 |
| #136 | -0.53543 | 0.94439 | -0.44363 | -0.53876 | -0.04843 | -0.41016 |
| #137 | -0.51182 | 1.02964 | -0.44534 | -0.53919 | -0.0548 | -0.4115 |
| #138 | -0.43666 | 2.40159 | -0.00441 | -0.23827 | 0.11909 | -0.00146 |
| #139 | -0.52663 | 1.11569 | -0.73117 | -0.57887 | -0.21745 | -0.66306 |
| #140 | -0.51874 | 1.43407 | 0.07977 | -0.1818 | 0.10513 | 0.06028 |
| #141 | -0.45886 | 1.62807 | -0.44386 | -0.26506 | -0.08784 | -0.39622 |
| #142 | -0.51141 | 1.1323 | -0.67067 | -0.57435 | -0.18035 | -0.60866 |
| #143 | -0.50904 | 0.92173 | -0.52647 | -0.56264 | -0.09968 | -0.48369 |
| #144 | -0.45781 | 1.44557 | -0.42015 | -0.47228 | -0.01855 | -0.38012 |
| #145 | -0.46621 | 1.15047 | -0.23886 | -0.412 | 0.04619 | -0.21487 |
| #146 | -0.43896 | 1.6862 | -0.33036 | -0.16101 | -0.07975 | -0.29442 |
| #147 | -0.47398 | 1.50734 | 0.15876 | -0.14875 | 0.14186 | 0.12943 |
| #148 | -0.43949 | 1.34689 | 0.34863 | -0.01444 | 0.22627 | 0.29528 |
| #149 | -0.46189 | 1.6101 | -0.82068 | -0.42122 | -0.32256 | -0.73303 |
| #150 | -0.45523 | 1.5337 | 0.28948 | -0.04453 | 0.19144 | 0.25207 |
| #151 | -0.46207 | 1.62752 | 0.27874 | -0.08879 | 0.20709 | 0.23876 |
| #152 | -0.4463 | 0.95063 | -0.25235 | -0.45914 | 0.03256 | -0.24022 |
| #153 | -0.41256 | 1.55632 | 0.23536 | -0.09993 | 0.16916 | 0.20184 |
| #154 | -0.45404 | 1.42937 | 0.23881 | -0.08849 | 0.17272 | 0.20786 |
| #155 | -0.4177 | 1.3845 | 0.30564 | -0.08319 | 0.22077 | 0.26115 |
| #156 | -0.45459 | 0.99947 | -0.68714 | -0.56672 | -0.19241 | -0.62389 |
| #157 | -0.43038 | 1.47678 | -0.76594 | -0.33618 | -0.28952 | -0.67441 |
| #158 | -0.36687 | 0.01276 | -0.32159 | -0.12865 | -0.13789 | -0.28691 |
| #159 | -0.39026 | 1.63222 | -0.41126 | -0.13204 | -0.14317 | -0.36453 |
| #160 | -0.4436 | 1.02837 | -0.64018 | -0.55559 | -0.13706 | -0.58075 |
| #161 | -0.43115 | 1.55275 | 0.39612 | 0.05392 | 0.21951 | 0.33535 |
| #162 | -0.41355 | 0.94584 | -0.51943 | -0.55969 | -0.08701 | -0.47788 |
| #163 | -0.43939 | 1.15748 | -0.70251 | -0.56575 | -0.20486 | -0.63515 |
| #164 | -0.42024 | 1.0208 | -0.60068 | -0.57641 | -0.14697 | -0.55011 |
| #165 | -0.37909 | 1.39566 | 0.29056 | -0.08194 | 0.22549 | 0.2512 |
| #166 | -0.43833 | 0.05216 | 0.69341 | 0.25073 | 0.31103 | 0.60951 |
| #167 | -0.39053 | 0.95512 | -0.42227 | -0.53272 | -0.04503 | -0.39193 |
| #168 | -0.3123 | 2.1407 | 0.25562 | -0.16618 | 0.26744 | 0.22492 |
| #169 | -0.34025 | 1.22538 | -0.37896 | -0.4869 | 0.01179 | -0.34543 |
| #170 | -0.32026 | 2.22256 | 0.24187 | -0.13357 | 0.24231 | 0.21717 |
| #171 | -0.36651 | 1.41835 | 0.30992 | -0.07273 | 0.22208 | 0.27027 |
| #172 | -0.29295 | 2.62759 | 0.10747 | -0.19966 | 0.19154 | 0.09924 |
| #173 | -0.36076 | 1.10659 | -0.64592 | -0.53859 | -0.16965 | -0.58056 |
| #174 | -0.29467 | 2.06437 | 0.11662 | -0.24319 | 0.21556 | 0.1028 |
| #175 | -0.35835 | 1.11466 | -0.71995 | -0.53069 | -0.21918 | -0.64467 |
| #176 | -0.33442 | 0.96424 | -0.65006 | -0.56982 | -0.17081 | -0.59041 |
| #177 | -0.34408 | 1.0186 | -0.60874 | -0.57673 | -0.14455 | -0.55711 |
| #178 | -0.25574 | 2.08312 | 0.0292 | -0.21513 | 0.13465 | 0.02908 |
| #179 | -0.34374 | 1.07138 | -0.69271 | -0.53454 | -0.19287 | -0.62145 |
| #180 | -0.31868 | 1.43757 | 0.12686 | -0.1786 | 0.14385 | 0.10464 |
| #181 | -0.4308 | 0.02048 | -0.5739 | -0.32604 | -0.24068 | -0.51635 |
| #182 | -0.28478 | 1.50142 | 0.26868 | -0.11216 | 0.21751 | 0.23088 |
| #183 | -0.27621 | 1.37028 | 0.12727 | -0.18296 | 0.14529 | 0.10554 |
| #184 | -0.32083 | 1.04867 | -0.69567 | -0.56594 | -0.19516 | -0.63032 |
| #185 | -0.29586 | 0.98362 | -0.67418 | -0.5435 | -0.1798 | -0.60581 |
| #186 | -0.29678 | 0.99223 | -0.73827 | -0.51429 | -0.2223 | -0.65957 |
| #187 | -0.23443 | 1.33338 | 0.58521 | 0.0638 | 0.36344 | 0.51651 |
| #188 | -0.26629 | 1.4668 | 0.27086 | -0.11238 | 0.20972 | 0.23052 |
| #189 | -0.30127 | 1.12518 | -0.64152 | -0.55494 | -0.15347 | -0.58089 |
| #190 | -0.24167 | 1.35883 | 0.36601 | -0.05938 | 0.26567 | 0.31942 |
| #191 | 0.074984 | 0.09265 | -0.05793 | -0.00731 | -0.01413 | -0.05965 |
| #192 | -0.24649 | 1.30963 | 0.29478 | -0.04147 | 0.20085 | 0.24392 |
| #193 | -0.30286 | 0.00475 | 0.14783 | -0.10765 | 0.06129 | 0.11751 |
| #194 | -0.28277 | 1.17315 | -0.66455 | -0.56154 | -0.17796 | -0.60085 |
| #195 | -0.2491 | 1.37159 | 0.38817 | -0.03246 | 0.26343 | 0.33532 |
| #196 | -0.27257 | 1.1463 | -0.64929 | -0.56205 | -0.17028 | -0.58909 |
| #197 | -0.19215 | 2.65421 | 0.16373 | -0.15497 | 0.19648 | 0.14775 |
| #198 | -0.28567 | 1.21709 | -0.7822 | -0.53985 | -0.25878 | -0.70402 |
| #199 | -0.28003 | 1.03621 | -0.67412 | -0.53549 | -0.18173 | -0.60714 |
| #200 | -0.28202 | 1.12999 | -0.71077 | -0.55619 | -0.20226 | -0.64296 |
| #201 | -0.23609 | 1.02386 | -0.367 | -0.5234 | 0.01466 | -0.34368 |
| #202 | -0.26753 | 1.01952 | -0.64283 | -0.56033 | -0.15287 | -0.58352 |
| #203 | -0.2513 | 1.03038 | -0.69821 | -0.56474 | -0.20608 | -0.63206 |
| #204 | -0.1993 | 1.68433 | -0.35547 | -0.17018 | -0.06911 | -0.31663 |
| #205 | -0.16904 | 2.55474 | 0.11415 | -0.16949 | 0.17695 | 0.10426 |
| #206 | -0.19044 | 1.36422 | 0.39992 | -0.01548 | 0.26139 | 0.34393 |
| #207 | -0.12997 | 1.21162 | 0.39562 | -0.06756 | 0.30036 | 0.34296 |
| #208 | -0.18894 | 1.5273 | 0.35881 | -0.05862 | 0.24332 | 0.31029 |
| #209 | -0.20948 | 1.06659 | -0.74598 | -0.56165 | -0.22199 | -0.67556 |
| #210 | -0.16511 | 1.38469 | 0.38819 | -0.03361 | 0.26216 | 0.3346 |
| #211 | -0.20744 | 1.10427 | -0.7796 | -0.55375 | -0.25195 | -0.70304 |
| #212 | -0.19721 | 1.67748 | 0.22424 | -0.06939 | 0.15843 | 0.19209 |
| #213 | -0.13532 | 0.87787 | -0.29996 | -0.49946 | 0.04856 | -0.28438 |
| #214 | -0.08001 | 1.51081 | -0.16723 | -0.31756 | 0.05961 | -0.14477 |
| #215 | -0.01451 | 1.57516 | 0.68392 | 0.12703 | 0.40592 | 0.5928 |
| #216 | -0.17503 | 1.11281 | -0.7847 | -0.5278 | -0.26307 | -0.70329 |
| #217 | -0.05105 | 1.20004 | 0.45143 | -0.03603 | 0.33851 | 0.39681 |
| #218 | -0.13032 | 1.15258 | -0.24298 | -0.40531 | -0.00033 | -0.23852 |
| #219 | -0.12756 | 0.99876 | -0.28042 | -0.4893 | 0.0472 | -0.26955 |
| #220 | -0.0448 | 2.19619 | 0.01874 | -0.19682 | 0.11986 | 0.0216 |
| #221 | -0.09306 | 1.38068 | 0.37158 | -0.04555 | 0.25554 | 0.31888 |
| #222 | -0.07371 | 1.23415 | -0.34919 | -0.47477 | 0.02704 | -0.31945 |
| #223 | -0.06577 | 1.379 | 0.26083 | -0.13598 | 0.23427 | 0.22425 |
| #224 | -0.08468 | 1.04769 | -0.65587 | -0.58589 | -0.17218 | -0.59855 |
| #225 | -0.08497 | 1.04157 | -0.46181 | -0.52194 | -0.03183 | -0.42106 |
| #226 | -0.11435 | 1.07617 | -0.71244 | -0.51951 | -0.20977 | -0.63711 |
| #227 | -0.02212 | 2.1402 | -0.29941 | -0.08213 | -0.09228 | -0.26609 |
| #228 | 0.11573 | 0.7603 | 0.53784 | 0.40537 | 0.18223 | 0.47471 |
| #229 | -0.07963 | 1.15165 | -0.4424 | -0.49592 | -0.01965 | -0.40071 |
| #230 | -0.1069 | 1.42764 | 0.24484 | -0.02785 | 0.15382 | 0.19897 |
| #231 | 0.011103 | 2.22602 | 0.33562 | 0.47895 | 0.04823 | 0.30831 |
| #232 | -0.11801 | 1.0795 | -0.73679 | -0.55294 | -0.22148 | -0.66669 |
| #233 | -0.04365 | 1.36587 | 0.26374 | -0.12907 | 0.22027 | 0.22399 |
| #234 | -0.05657 | 1.3941 | 0.36819 | -0.05783 | 0.25916 | 0.31655 |
| #235 | -0.08892 | 1.11112 | -0.78142 | -0.5193 | -0.25347 | -0.70341 |
| #236 | -0.0034 | 1.06836 | 0.50828 | 0.01891 | 0.34106 | 0.44465 |
| #237 | 0.021201 | 2.36105 | 0.17471 | 0.32867 | 0.00641 | 0.16082 |
| #238 | -0.03444 | 1.01323 | -0.43169 | -0.53815 | -0.00261 | -0.39835 |
| #239 | 0.024449 | 2.92343 | 0.29328 | -0.08066 | 0.25408 | 0.26304 |
| #240 | -0.02202 | 1.62771 | 0.12928 | -0.14129 | 0.11986 | 0.10666 |
| #241 | -0.00257 | 1.99125 | -0.33468 | -0.14067 | -0.07278 | -0.29776 |
| #242 | -0.01812 | 54.79931 | -0.0626 | -0.00608 | -0.01976 | -0.06238 |
| #243 | 0.00014 | 11.42423 | -0.00709 | -0.12666 | 0.08842 | -0.01247 |
| #244 | -0.03834 | 1.08877 | -0.75211 | -0.53544 | -0.23661 | -0.67757 |
| #245 | -0.01949 | 1.61445 | -0.54503 | -0.22975 | -0.17895 | -0.47996 |
| #246 | 0.041249 | 0.13902 | -0.4099 | -0.0175 | -0.27952 | -0.35696 |
| #247 | 0.006201 | 1.413 | 0.0647 | -0.21047 | 0.1072 | 0.04609 |
| #248 | 0.040287 | 0.01051 | 0.3084 | 0.10287 | 0.1627 | 0.27798 |
| #249 | 0.073466 | 2.45922 | 0.06785 | -0.15874 | 0.14727 | 0.06499 |
| #250 | 0.017387 | 1.40041 | 0.32646 | -0.07561 | 0.23657 | 0.27912 |
| #251 | 0.080848 | 1.27016 | 0.42675 | -0.05321 | 0.31332 | 0.37006 |
| #252 | 0.120847 | 1.26697 | 0.34041 | 0.03928 | 0.19199 | 0.28722 |
| #253 | 0.058537 | 1.68281 | -0.3053 | -0.40727 | 0.03795 | -0.27313 |
| #254 | 0.056362 | 1.41637 | 0.33736 | -0.06754 | 0.2365 | 0.28885 |
| #255 | 0.054456 | 1.61546 | 0.20839 | -0.10532 | 0.15679 | 0.17763 |
| #256 | 0.041843 | 0.95677 | -0.58403 | -0.56641 | -0.1296 | -0.53336 |
| #257 | 0.064956 | 1.68102 | 0.1455 | -0.13644 | 0.12949 | 0.12053 |
| #258 | 0.103011 | 1.39812 | 0.53283 | 0.06449 | 0.31588 | 0.46596 |
| #259 | 0.122502 | 0.88639 | 0.05349 | -0.33644 | 0.23283 | 0.03247 |
| #260 | 0.151313 | 1.76258 | -0.19051 | -0.13543 | 0.0125 | -0.17452 |
| #261 | 0.16803 | 2.21065 | -0.07202 | -0.1805 | 0.05521 | -0.05811 |
| #262 | 0.108078 | 1.527 | 0.14431 | -0.15586 | 0.13391 | 0.11917 |
| #263 | 0.111068 | 1.42359 | 0.38237 | -0.03927 | 0.26037 | 0.32954 |
| #264 | 0.081339 | 1.06463 | -0.76405 | -0.54962 | -0.24613 | -0.68816 |
| #265 | 0.176676 | 2.4052 | -0.16958 | 0.00502 | -0.03073 | -0.14997 |
| #266 | 0.1896 | 4.09163 | -0.00952 | 0.14207 | -0.00245 | -0.00191 |
| #267 | 0.137521 | 1.37823 | 0.34252 | -0.04892 | 0.23734 | 0.29065 |
| #268 | 0.155126 | 1.55203 | 0.35855 | 0.0173 | 0.2038 | 0.29972 |
| #269 | 0.140289 | 1.06754 | -0.24676 | -0.46264 | 0.1051 | -0.23447 |
| #270 | 0.173334 | 1.8175 | -0.4151 | -0.14143 | -0.13265 | -0.3654 |
| #271 | 0.20276 | 2.84456 | -0.27865 | -0.21892 | -0.00644 | -0.24578 |
| #272 | 0.228513 | 0.83179 | 0.06397 | -0.33261 | 0.247 | 0.04301 |
| #273 | 0.140405 | 0.00457 | 0.11934 | -0.07487 | -0.01356 | 0.08855 |
| #274 | 0.243513 | 1.77689 | -0.03588 | -0.33272 | 0.18286 | -0.03709 |
| #275 | 0.216436 | 1.22547 | 0.16157 | -0.11207 | 0.14851 | 0.12279 |
| #276 | 0.242949 | 7.55518 | -0.00696 | 0.15052 | -0.00254 | 0.00301 |
| #277 | 0.271082 | 3.64138 | 0.18879 | -0.14011 | 0.22549 | 0.17148 |
| #278 | 0.251422 | 8.40826 | -0.13574 | -0.01901 | -0.02492 | -0.11842 |
| #279 | 0.228954 | 1.56095 | 0.18807 | -0.14314 | 0.15473 | 0.15802 |
| #280 | 0.271731 | 1.25225 | -0.03016 | -0.36672 | 0.19838 | -0.03728 |
| #281 | 0.297887 | 3.69306 | 0.18317 | 0.0861 | 0.14667 | 0.15819 |
| #282 | 0.202433 | 0.02647 | -0.25222 | -0.03572 | -0.2371 | -0.23737 |
| #283 | 0.243574 | 0.91284 | -0.40762 | -0.53293 | 0.00011 | -0.37973 |
| #284 | 0.263818 | 2.02296 | -0.39343 | -0.30701 | -0.06064 | -0.34792 |
| #285 | 0.29596 | 1.62619 | -0.26399 | -0.19998 | 0.00761 | -0.238 |
| #286 | 0.281611 | 2.36045 | -0.28548 | -0.09555 | -0.06561 | -0.25125 |
| #287 | 0.311107 | 1.38199 | 0.34064 | -0.00059 | 0.20713 | 0.28221 |
| #288 | 0.333138 | 0.8176 | 0.00946 | -0.36051 | 0.22376 | -0.00578 |
| #289 | 0.317853 | 1.82933 | -0.23169 | -0.36374 | 0.0753 | -0.2115 |
| #290 | 0.335661 | 3.35064 | 0.01178 | -0.19951 | 0.172 | 0.01152 |
| #291 | 0.32645 | 1.68539 | 0.08844 | -0.14651 | 0.09277 | 0.07128 |
| #292 | 0.306307 | 1.52807 | 0.42346 | 0.0362 | 0.25608 | 0.3621 |
| #293 | 0.301109 | 1.45903 | 0.40517 | -0.02239 | 0.27198 | 0.34695 |
| #294 | 0.319596 | 2.60711 | -0.35494 | -0.17551 | -0.08622 | -0.31184 |
| #295 | 0.367425 | 0.00516 | 0.38824 | -0.02642 | 0.25317 | 0.33766 |
| #296 | 0.320776 | 4.06617 | -0.3615 | -0.11899 | -0.12181 | -0.3153 |
| #297 | 0.403138 | 2.88917 | 0.08727 | 0.1892 | 0.04783 | 0.084 |
| #298 | 0.32312 | 1.74315 | -0.49273 | -0.25233 | -0.15303 | -0.43423 |
| #299 | 0.369375 | 9.67105 | 0.14809 | 0.00796 | 0.14647 | 0.13024 |
| #300 | 0.35443 | 1.45998 | 0.32674 | -0.0878 | 0.24575 | 0.27756 |
| #301 | 0.416424 | 3.08856 | -0.11142 | -0.04483 | 0.03644 | -0.09663 |
| #302 | 0.393172 | 2.09395 | -0.42536 | -0.19194 | -0.12465 | -0.37312 |
| #303 | 0.455611 | 3.0309 | 0.07695 | 0.07912 | 0.10384 | 0.06771 |
| #304 | 0.425418 | 4.40369 | -0.1971 | 0.03235 | -0.08337 | -0.17222 |
| #305 | 0.438318 | 1.98021 | -0.25207 | -0.32551 | 0.04565 | -0.2277 |
| #306 | 0.454324 | 0.89667 | 0.06682 | -0.3273 | 0.2513 | 0.04617 |
| #307 | 0.467786 | 1.1691 | 0.05873 | -0.30831 | 0.24542 | 0.04313 |
| #308 | 0.366637 | 1.34318 | 0.09588 | -0.23854 | 0.16334 | 0.06965 |
| #309 | 0.449944 | 2.68722 | -0.19362 | -0.23781 | 0.05639 | -0.17488 |
| #310 | 0.493345 | 1.00162 | 0.0878 | -0.31726 | 0.25877 | 0.06644 |
| #311 | 0.393842 | 1.77677 | -0.46703 | -0.22197 | -0.1423 | -0.40942 |
| #312 | 0.447951 | 2.42193 | -0.35121 | -0.2645 | -0.04656 | -0.31139 |
| #313 | 0.492052 | 0.83657 | 0.89054 | 0.36302 | 0.38015 | 0.7895 |
| #314 | 0.409661 | 1.55047 | -0.58668 | -0.33262 | -0.17636 | -0.51856 |
| #315 | 0.388289 | 1.60971 | 0.27755 | 0.02474 | 0.15788 | 0.23161 |
| #316 | 0.39476 | 1.12705 | -0.80804 | -0.45866 | -0.26729 | -0.72284 |
| #317 | 0.50961 | 3.59957 | -0.10208 | -0.03919 | 0.03076 | -0.08664 |
| #318 | 0.504706 | 2.01183 | -0.2315 | -0.29279 | 0.05532 | -0.20872 |
| #319 | 0.462631 | 2.88728 | -0.33106 | -0.126 | -0.09074 | -0.28766 |
| #320 | 0.510681 | 2.43826 | -0.25896 | -0.28369 | 0.0235 | -0.23134 |
| #321 | 0.48536 | 1.5439 | 0.50757 | 0.03093 | 0.31902 | 0.44433 |
| #322 | 0.497324 | 2.05912 | -0.36129 | -0.25697 | -0.05155 | -0.31969 |
| #323 | 0.522056 | 1.4735 | 0.6307 | 0.1537 | 0.337 | 0.55107 |
| #324 | 0.510793 | 2.77895 | -0.32033 | -0.15356 | -0.08289 | -0.27994 |
| #325 | 0.649045 | 1.21453 | 0.30133 | 0.10599 | 0.16587 | 0.2586 |
| #326 | 0.529977 | 1.28596 | 0.31973 | -0.03923 | 0.2264 | 0.26473 |
| #327 | 0.509564 | 1.44298 | 0.24504 | -0.14103 | 0.21063 | 0.20733 |
| #328 | 0.48891 | 0.13507 | -0.03455 | -0.20475 | 0.02844 | -0.03851 |
| #329 | 0.545457 | 1.46939 | 0.58322 | 0.10423 | 0.32226 | 0.50888 |
| #330 | 0.530015 | 2.44188 | -0.34529 | -0.17328 | -0.08063 | -0.30326 |
| #331 | 0.581838 | 4.81246 | 0.14649 | 0.04271 | 0.15894 | 0.12764 |
| #332 | 0.583697 | 4.11511 | 0.05664 | 0.13141 | 0.05337 | 0.05304 |
| #333 | 0.512165 | 1.59563 | 0.43888 | 0.05912 | 0.24922 | 0.37232 |
| #334 | 0.5934 | 3.31512 | 0.33812 | -0.09907 | 0.31931 | 0.30077 |
| #335 | 0.599016 | 3.0387 | 0.14304 | -0.15134 | 0.23165 | 0.12848 |
| #336 | 0.216016 | 0.34059 | 0.49104 | 0.19503 | 0.25882 | 0.45899 |
| #337 | 0.55341 | 1.89118 | -0.14591 | 0.03386 | -0.05598 | -0.11714 |
| #338 | 0.613416 | 1.83266 | -0.09128 | -0.08635 | 0.06189 | -0.08644 |
| #339 | 0.61123 | 5.95983 | 0.22637 | 0.17073 | 0.14343 | 0.20325 |
| #340 | 0.611036 | 4.47037 | 0.05412 | -0.05766 | 0.13798 | 0.04748 |
| #341 | 0.616504 | 5.3154 | 0.11224 | 0.10646 | 0.10535 | 0.10079 |
| #342 | 0.620803 | 4.78073 | 0.12918 | 0.05449 | 0.14705 | 0.1156 |
| #343 | 0.608711 | 5.26337 | 0.72302 | 0.25745 | 0.40587 | 0.64203 |
| #344 | 0.616336 | 4.06236 | -0.00095 | -0.05522 | 0.11884 | 0.00138 |
| #345 | 0.581254 | 2.42138 | -0.34571 | -0.16679 | -0.0886 | -0.30143 |
| #346 | 0.611226 | 2.62006 | -0.11629 | -0.05411 | 0.03802 | -0.10017 |
| #347 | 0.632142 | 4.32613 | 0.14697 | 0.08772 | 0.13688 | 0.13183 |
| #348 | 0.62906 | 4.34244 | -0.01199 | 0.06568 | 0.04814 | -0.00483 |
| #349 | 0.625441 | 2.68694 | -0.09875 | -0.04136 | 0.04562 | -0.08532 |
| #350 | 0.612123 | 2.20231 | -0.24445 | -0.15533 | -0.00551 | -0.21522 |
| #351 | 0.629679 | 1.65047 | 0.38631 | -0.007 | 0.23719 | 0.33438 |
| #352 | 0.555798 | 1.85098 | -0.44269 | -0.18848 | -0.13898 | -0.39181 |
| #353 | 0.634142 | 2.53702 | -0.30009 | -0.14788 | -0.05211 | -0.26302 |
| #354 | 0.639167 | 2.79667 | -0.14876 | -0.08673 | 0.03041 | -0.12861 |
| #355 | 0.645472 | 2.18315 | -0.26569 | -0.16395 | -0.01615 | -0.23323 |
| #356 | 0.673991 | 1.40463 | 0.67957 | 0.17833 | 0.36409 | 0.59845 |
| #357 | 0.608016 | 1.74366 | -0.49885 | -0.32751 | -0.13091 | -0.44269 |
| #358 | 0.5887 | 0.00908 | 0.22242 | 0.13934 | 0.11815 | 0.20382 |
| #359 | 0.695056 | 0.97855 | 0.28837 | -0.18857 | 0.31949 | 0.24407 |
| #360 | 0.685073 | 4.87436 | 0.11404 | -0.02648 | 0.16769 | 0.10049 |
| #361 | 0.683479 | 2.92172 | -0.08391 | -0.10862 | 0.0803 | -0.07285 |
| #362 | 0.686097 | 2.14553 | -0.20412 | -0.28032 | 0.06651 | -0.18265 |
| #363 | 0.690381 | 2.32644 | -0.16163 | -0.24732 | 0.06999 | -0.14151 |
| #364 | 0.665919 | 2.01401 | -0.25467 | -0.07378 | -0.04549 | -0.22421 |
| #365 | 0.688476 | 2.46482 | -0.25566 | -0.16412 | -0.01424 | -0.22411 |
| #366 | 0.713681 | 3.90543 | 0.12152 | 0.0071 | 0.16353 | 0.10448 |
| #367 | 0.703687 | 2.73281 | -0.21682 | -0.11268 | -0.01255 | -0.18891 |
| #368 | 0.749216 | 1.54236 | 0.48349 | 0.21212 | 0.22136 | 0.42003 |
| #369 | 0.719841 | 3.80265 | 0.03267 | 0.06162 | 0.07672 | 0.03298 |
| #370 | 0.725613 | 3.99295 | -0.11381 | -0.02809 | 0.01266 | -0.09572 |
| #371 | 0.72047 | 2.82701 | -0.27758 | -0.14871 | -0.05118 | -0.24347 |
| #372 | 0.758424 | 4.28655 | 0.26739 | -0.04276 | 0.25512 | 0.24269 |
| #373 | 0.764544 | 5.00831 | -0.01531 | 0.03454 | 0.05454 | -0.00793 |
| #374 | 0.766743 | 0.00541 | 0.62335 | 0.19709 | 0.36046 | 0.55788 |
| #375 | 0.794735 | 4.33429 | -0.08253 | -0.00916 | 0.0293 | -0.06853 |
| #376 | 0.803751 | 3.70912 | 0.10443 | -0.1201 | 0.20457 | 0.09034 |
| #377 | 0.69613 | 1.51322 | 0.29349 | -0.10208 | 0.22617 | 0.24015 |
| #378 | 0.797297 | 1.94825 | 0.13887 | -0.17322 | 0.26229 | 0.11588 |
| #379 | 0.832621 | 1.40743 | 0.21511 | -0.23322 | 0.3115 | 0.18285 |
| #380 | 0.81971 | 2.29591 | -0.01978 | -0.21825 | 0.15967 | -0.02321 |
| #381 | 0.822111 | 4.23923 | -0.0942 | -0.02352 | 0.02919 | -0.07833 |
| #382 | 0.783231 | 1.50587 | 0.57567 | 0.08146 | 0.32845 | 0.50331 |
| #383 | 0.838334 | 3.75805 | 0.09536 | -0.01057 | 0.15842 | 0.08246 |
| #384 | 0.85219 | 3.88448 | 0.13717 | 0.18357 | 0.08248 | 0.12298 |
| #385 | 0.857138 | 1.89999 | 0.17095 | -0.19979 | 0.28526 | 0.14518 |
| #386 | 0.649956 | 0.49511 | -0.61973 | -0.13165 | -0.36733 | -0.55307 |
| #387 | 0.869066 | 1.82446 | 0.32776 | -0.13153 | 0.3511 | 0.28468 |
| #388 | 0.84517 | 2.98543 | -0.20933 | -0.07772 | -0.0197 | -0.1813 |
| #389 | 0.709012 | 0.03564 | -0.85058 | -0.34387 | -0.41204 | -0.74568 |
| #390 | 1.011813 | 1.1902 | 0.47664 | 0.24763 | 0.24099 | 0.42145 |
| #391 | 0.879106 | 6.46621 | 0.64054 | 0.44053 | 0.24093 | 0.56495 |
| #392 | 0.859469 | 1.91636 | -0.18914 | -0.11501 | 0.0207 | -0.17033 |
| #393 | 0.889807 | 1.30775 | 0.19604 | 0.02168 | 0.12909 | 0.16109 |
| #394 | 0.866624 | 5.40763 | 0.1837 | 0.03138 | 0.19368 | 0.15691 |
| #395 | 0.914214 | 1.49365 | 0.36507 | -0.13717 | 0.36944 | 0.31797 |
| #396 | 0.925095 | 1.32527 | 0.26336 | 0.04668 | 0.15306 | 0.22196 |
| #397 | 0.840466 | 0.2693 | 0.36067 | 0.24185 | 0.17157 | 0.33512 |
| #398 | 0.905653 | 4.62406 | 0.23147 | 0.08589 | 0.1978 | 0.2024 |
| #399 | 0.699792 | 0.05198 | -0.41229 | -0.38406 | -0.11245 | -0.37121 |
| #400 | 1.00394 | 1.18966 | 0.23105 | 0.03992 | 0.17828 | 0.19778 |
| #401 | 0.931294 | 2.4327 | 0.48769 | -0.03586 | 0.40604 | 0.42797 |
| #402 | 1.038902 | 1.21532 | 0.48026 | 0.22352 | 0.24664 | 0.42308 |
| #403 | 0.760969 | 1.32791 | -0.80057 | -0.422 | -0.27632 | -0.70917 |
| #404 | 0.870252 | 2.77205 | -0.1422 | 0.02587 | -0.03736 | -0.12092 |
| #405 | 0.872592 | 0.03629 | -0.09981 | -0.20725 | 0.04542 | -0.08737 |
| #406 | 0.957937 | 4.08073 | 0.20109 | 0.14744 | 0.15154 | 0.17782 |
| #407 | 1.14914 | 0.00948 | 0.73111 | 0.18368 | 0.36664 | 0.63843 |
| #408 | 0.969058 | 2.91736 | 0.34736 | -0.0559 | 0.33676 | 0.30239 |
| #409 | 0.970259 | 4.33849 | 0.23197 | 0.09213 | 0.19349 | 0.20283 |
| #410 | 0.634261 | 0.00598 | 0.09927 | -0.02784 | -0.00429 | 0.08033 |
| #411 | 0.99274 | 1.91545 | 0.86994 | 0.48787 | 0.3669 | 0.7847 |
| #412 | 0.976444 | 2.11689 | 0.28487 | -0.14363 | 0.33063 | 0.24706 |
| #413 | 0.980297 | 1.2249 | 0.20669 | -0.21182 | 0.30971 | 0.17461 |
| #414 | 0.99068 | 5.57494 | 0.30513 | 0.25286 | 0.1563 | 0.27202 |
| #415 | 0.992207 | 3.42369 | 0.13921 | 0.07033 | 0.14489 | 0.12162 |
| #416 | 0.894623 | 1.64786 | -0.45728 | -0.23238 | -0.13 | -0.40213 |
| #417 | 0.964281 | 1.19356 | 0.11979 | -0.26353 | 0.22113 | 0.09312 |
| #418 | 1.390137 | 0.02156 | -0.51299 | -0.0027 | -0.34331 | -0.44428 |
| #419 | 1.013866 | 1.37213 | 0.54128 | -0.03153 | 0.4202 | 0.47499 |
| #420 | 1.058476 | 1.27831 | -0.7956 | -0.27531 | -0.35828 | -0.6997 |
| #421 | 1.015607 | 1.40127 | 0.36602 | -0.13635 | 0.37078 | 0.3176 |
| #422 | 1.004323 | 3.16886 | 0.03275 | -0.04417 | 0.1396 | 0.02766 |
| #423 | 1.010966 | 3.13278 | 0.23233 | 0.12041 | 0.16258 | 0.20461 |
| #424 | 1.01471 | 3.16953 | 0.13331 | 0.05308 | 0.15201 | 0.11526 |
| #425 | 1.052664 | 3.45436 | 0.88448 | 0.35607 | 0.43429 | 0.78079 |
| #426 | 0.947221 | 1.52727 | -0.21425 | -0.05929 | -0.07661 | -0.1835 |
| #427 | 1.044749 | 1.97914 | 0.25418 | -0.20168 | 0.31536 | 0.2177 |
| #428 | 1.030488 | 3.7334 | 0.4385 | 0.17029 | 0.26321 | 0.38403 |
| #429 | 1.02683 | 3.93277 | 0.20577 | 0.13242 | 0.16088 | 0.18078 |
| #430 | 1.02821 | 1.40489 | 0.31404 | -0.15429 | 0.3516 | 0.2716 |
| #431 | 1.037555 | 3.71805 | 0.18801 | 0.13765 | 0.14086 | 0.16546 |
| #432 | 1.035309 | 0.97128 | 0.17216 | -0.25905 | 0.30028 | 0.14306 |
| #433 | 0.861323 | 0.03585 | 0.50286 | 0.14243 | 0.26035 | 0.45532 |
| #434 | 1.10883 | 1.23077 | 0.34304 | 0.10926 | 0.19677 | 0.29674 |
| #435 | 0.881276 | 0.00701 | -0.10838 | -0.09991 | -0.02315 | -0.10373 |
| #436 | 1.115642 | 2.45714 | 0.4071 | 0.13388 | 0.25743 | 0.3538 |
| #437 | 1.096939 | 2.94273 | 0.2317 | 0.11469 | 0.17808 | 0.20447 |
| #438 | 1.109652 | 5.16891 | 0.51591 | 0.31241 | 0.23516 | 0.45516 |
| #439 | 1.11398 | 2.6499 | 0.4646 | -0.02596 | 0.38997 | 0.40661 |
| #440 | 1.304638 | 0.03028 | 0.74718 | 0.25842 | 0.30999 | 0.6549 |
| #441 | 1.120053 | 5.16795 | 0.65209 | 0.27091 | 0.3221 | 0.5748 |
| #442 | 1.11816 | 3.9566 | 0.31829 | 0.1075 | 0.24375 | 0.27551 |
| #443 | 0.79236 | 0.01083 | 0.4876 | 0.34366 | 0.08566 | 0.43227 |
| #444 | 1.143966 | 1.13138 | 0.89702 | 0.40001 | 0.4412 | 0.80279 |
| #445 | 1.125275 | 1.43748 | 0.41094 | -0.09678 | 0.38469 | 0.35764 |
| #446 | 1.145842 | 3.74576 | 0.5152 | 0.21596 | 0.28748 | 0.45381 |
| #447 | 1.088208 | 1.07638 | -0.2104 | 0.01301 | -0.10448 | -0.18103 |
| #448 | 1.152329 | 1.66052 | 0.49422 | -0.02137 | 0.40394 | 0.43233 |
| #449 | 1.153433 | 1.56189 | 0.39839 | -0.09862 | 0.38167 | 0.34755 |
| #450 | 1.180186 | 3.56408 | 0.5037 | 0.34949 | 0.20933 | 0.45124 |
| #451 | 1.164945 | 2.01961 | 0.55057 | 0.01972 | 0.42011 | 0.483 |
| #452 | 0.966036 | 0.02212 | -0.81161 | -0.39379 | -0.36395 | -0.72202 |
| #453 | 1.193731 | 4.4551 | 0.60105 | 0.33204 | 0.26662 | 0.53397 |
| #454 | 1.191814 | 1.3867 | 0.44702 | -0.07939 | 0.39795 | 0.39017 |
| #455 | 1.175706 | 3.53628 | 0.49874 | 0.03437 | 0.38561 | 0.43627 |
| #456 | 1.215845 | 3.37104 | 0.41856 | 0.16923 | 0.24921 | 0.36749 |
| #457 | 1.216663 | 3.91737 | 0.48199 | 0.18411 | 0.28022 | 0.42077 |
| #458 | 1.234633 | 3.69173 | 0.58256 | 0.34747 | 0.2518 | 0.51639 |
| #459 | 1.275694 | 2.27995 | 0.84964 | 0.41867 | 0.39637 | 0.75737 |
| #460 | 1.326786 | 2.71209 | 0.49174 | 0.32336 | 0.21817 | 0.4355 |
| #461 | 1.313649 | 1.15431 | 0.45901 | 0.28977 | 0.17601 | 0.40508 |
| #462 | 1.313495 | 0.64698 | 0.8271 | 0.49597 | 0.35997 | 0.74681 |
| #463 | 1.327424 | 1.38733 | 0.46949 | 0.19346 | 0.2408 | 0.40822 |
| #464 | 1.334502 | 1.55095 | 0.41318 | 0.19372 | 0.19595 | 0.35811 |
| #465 | 1.271204 | 1.63463 | 0.36838 | 0.10584 | 0.18994 | 0.31521 |
| #466 | 1.314732 | 2.13447 | 0.60767 | 0.07486 | 0.42288 | 0.53227 |
| #467 | 1.344866 | 3.05349 | 0.55876 | 0.24272 | 0.30035 | 0.49624 |
| #468 | 1.384701 | 2.11507 | 0.37678 | 0.28003 | 0.17953 | 0.33706 |
| #469 | 1.299217 | 0.04415 | -0.87073 | -0.44615 | -0.36615 | -0.77044 |
| #470 | 1.387872 | 2.08918 | 0.8863 | 0.3112 | 0.47659 | 0.79058 |
| #471 | 1.410051 | 2.28648 | 0.4586 | 0.28466 | 0.22316 | 0.40926 |
| #472 | 1.401647 | 2.27346 | 0.6064 | 0.24007 | 0.32492 | 0.53601 |
| #473 | 1.397144 | 2.54455 | 0.89344 | 0.31783 | 0.47685 | 0.79611 |
| #474 | 1.492475 | 0.02442 | 0.62541 | 0.17775 | 0.27032 | 0.54192 |
| #475 | 1.435674 | 2.32478 | 0.76549 | 0.16378 | 0.47322 | 0.67608 |
| #476 | 1.482068 | 2.47311 | 0.62412 | 0.22275 | 0.34266 | 0.55047 |
| #477 | 1.499263 | 2.46185 | 0.64042 | 0.21657 | 0.35589 | 0.56542 |
| #478 | 1.496302 | 3.97502 | 0.7365 | 0.17719 | 0.44749 | 0.64895 |
| #479 | 1.554522 | 2.45888 | 0.63644 | 0.32889 | 0.29168 | 0.56615 |
| #480 | 1.2518 | 0.02947 | -0.76 | -0.36968 | -0.3783 | -0.68006 |
| #481 | 1.706623 | 1.14231 | 0.40265 | 0.30242 | 0.18763 | 0.36081 |
| #482 | 1.659529 | 0.19839 | 0.64313 | 0.09893 | 0.41947 | 0.56822 |
| #483 | 1.612124 | 14.32482 | 0.67515 | 0.19957 | 0.3257 | 0.60759 |
| #484 | 1.762399 | 2.46629 | 0.84884 | 0.26028 | 0.47204 | 0.75125 |
| #485 | 1.493937 | 0.04277 | -0.46027 | -0.30727 | -0.2062 | -0.42058 |
| #486 | 1.875567 | 0.20474 | 0.78498 | 0.33511 | 0.35317 | 0.69672 |
| #487 | 1.800546 | 0.3228 | 0.6077 | 0.18707 | 0.29751 | 0.53322 |
| #488 | 1.972766 | 1.77012 | 0.44356 | 0.10161 | 0.30163 | 0.38709 |
| #489 | 1.936641 | 0.00416 | 0.6593 | 0.10797 | 0.38115 | 0.57325 |
| #490 | 1.872525 | 0.04176 | -0.83458 | -0.35141 | -0.41006 | -0.73695 |
| #491 | 2.152826 | 2.53155 | 0.85521 | 0.26884 | 0.47048 | 0.75699 |
| #492 | 2.188821 | 1.27328 | 0.68574 | 0.17717 | 0.4018 | 0.60059 |
| #493 | 2.709568 | 0.01479 | 0.31812 | 0.15614 | 0.10464 | 0.277 |
| #494 | 2.172443 | 0.01399 | -0.48664 | -0.26534 | -0.16796 | -0.42889 |
| #495 | 2.212508 | 0.01982 | 0.2671 | 0.09672 | 0.11782 | 0.24443 |
| #496 | 2.459038 | 0.05788 | -0.66922 | -0.17821 | -0.35114 | -0.57747 |
| #497 | 2.941915 | 1.40229 | 0.79425 | 0.23331 | 0.46482 | 0.70821 |
| #498 | 2.517684 | 0.01099 | 0.48816 | 0.39074 | 0.17468 | 0.44956 |
| #499 | 2.833806 | 0.0348 | 0.18667 | 0.19015 | 0.04058 | 0.15931 |
| #500 | 2.837643 | 0.04521 | -0.62769 | -0.23138 | -0.26002 | -0.53802 |
| #501 | 3.437767 | 0.28281 | 0.65991 | 0.1765 | 0.35412 | 0.58031 |
| #502 | 3.647944 | 0.01973 | 0.85498 | 0.42922 | 0.37159 | 0.75784 |
| #503 | 4.085928 | 0.01085 | 0.39187 | 0.18016 | 0.10174 | 0.34195 |
| #504 | 3.81938 | 0.0097 | -0.11827 | 0.17838 | -0.08667 | -0.09165 |
| #505 | 4.044226 | 0.0474 | -0.1184 | -0.28852 | 0.03047 | -0.11104 |
| #506 | 4.047894 | 0.01052 | 0.08718 | 0.11035 | 0.06572 | 0.07652 |
| #507 | 6.578578 | 0.02937 | 0.85702 | 0.3846 | 0.41456 | 0.76133 |
| #508 | 6.045519 | 0.01614 | -0.68206 | -0.27462 | -0.27825 | -0.59563 |
| #509 | 6.305846 | 0.00717 | 0.75083 | 0.27325 | 0.41577 | 0.66871 |
| #510 | 8.043719 | 0.02003 | -0.82768 | -0.29451 | -0.40327 | -0.71617 |
| #511 | 9.670453 | 0.04383 | -0.24373 | -0.208 | -0.02748 | -0.19979 |

**Table S2 | Collected samples from each patient.**

| Patient ID (n=41) | N (n=7) | T (n=28) | M (n=27) |
| --- | --- | --- | --- |
| #0939155 |  | √ |  |
| #1763629 |  | √ | √ |
| #1809708 |  | √ |  |
| #1826545 |  | √ | √ |
| #1831631 |  |  | √ |
| #1832060 |  | √ | √ |
| #1833417 | √ |  |  |
| #1835207 |  | √ |  |
| #1835227 |  | √ | √ |
| #1837519 |  | √ | √ |
| #1841932 |  | √ | √ |
| #1850109 |  | √ | √ |
| #1850806 |  |  | √ |
| #1853111 |  | √ | √ |
| #1855139 |  | √ | √ |
| #1874576 |  | √ | √ |
| #1875931 | √ |  |  |
| #1877627 |  | √ | √ |
| #1883328 |  |  | √ |
| #1883971 |  |  | √ |
| #1886103 |  |  | √ |
| #1887896 |  | √ | √ |
| #1899673 |  | √ | √ |
| #1900985 | √ |  |  |
| #1901246 |  | √ | √ |
| #1905174 |  | √ | √ |
| #1907404 |  | √ |  |
| #1909357 |  |  | √ |
| #1931850 |  | √ | √ |
| #1936419 |  | √ |  |
| #1938711 |  |  | √ |
| #1944493 |  | √ |  |
| #1955514 |  | √ |  |
| #2038446 |  | √ | √ |
| #2044864 |  | √ | √ |
| #2047500 |  | √ | √ |
| #2075235 | √ |  |  |
| #2095674 | √ | √ |  |
| #2104514 | √ |  |  |
| #2121724 |  | √ | √ |
| #9593717 | √ |  |  |

**Table. S3 | Ground truths and model training labels defined for different resection cases.**

| First frozen section diagnosis | Permanent section diagnosis | Ground truth | Model training label | Number of patients |
| --- | --- | --- | --- | --- |
| Negative | Negative | Negative | Negative | 17 |
| Positive | Negative | Positive | Positive | 5 |
| Negative | Positive | Positive | Not used for training | 2 |
| Positive | Positive | Positive | Positive | 3 |

**Table S4 | Frozen and permanent section diagnoses, COMP#475 values, and MICLEAR-predicted patient-level positive probability of 27 patients with *M* samples.**

|  | First frozen section diagnosis | Permanent section diagnosis | COMP#475 maximum | Positive probability (%) |
| --- | --- | --- | --- | --- |
| NEG#1 | Negative | Negative^&&^ | 1.233 | 12.87 |
| NEG#2 | Negative | Negative | 1.471 | 50.34 |
| NEG#3 | Negative | Negative | 0.720 | 0.07 |
| NEG#4 | Negative | Negative | 0.561 | 1.86 |
| NEG#5 | Negative | Negative | 0.983 | 1.11 |
| NEG#6 | Negative | Negative | 1.480 | 0.80 |
| NEG#7 | Negative | Negative | 0.840 | 0.41 |
| NEG#8 | Negative | Negative | 2.239 | 18.73 |
| NEG#9 | Negative | Negative | 1.361 | 1.24 |
| NEG#10 | Negative | Negative | 1.241 | 27.96 |
| NEG#11 | Negative | Negative | 0.884 | 18.72 |
| NEG#12 | Negative | Negative | 1.040 | 2.00 |
| NEG#13 | Negative | Negative | 0.906 | 4.01 |
| NEG#14 | Negative | Negative | 0.633 | 13.76 |
| NEG#15 | Negative | Negative | 0.766 | 18.58 |
| NEG#16 | Negative | Negative | 1.459 | 28.24 |
| NEG#17 | Negative | Negative | 1.472 | 4.46 |
| POS#1* | Positive | Positive | 1.784 | 94.12 |
| POS#2* | Positive | Positive | 3.363 | 99.99 |
| POS#3* | Positive | Negative | 2.124 | 99.96 |
| POS#4* | Positive | Negative | 1.628 | 88.47 |
| POS#5* | Positive | Negative | 1.479 | 59.10 |
| POS#6* | Positive | Negative | 2.191 | 99.86 |
| POS#7** | Negative | Positive | 0.479 | 0.03 |
| POS#8* | Positive | Positive | 1.094 | 8.26 |
| POS#9* | Positive | Negative | 3.446 | 99.99 |
| POS#10** | Negative | Positive | 2.348 | 99.99 |

* The permanent section was sampled at a location different from the first frozen section and the first frozen section diagnosis served as the ground truth.

** The permanent section was sampled at the same location as the first frozen section, but their diagnoses were inconsistent. The permanent section result served as the ground truth. The sample was not used for model training.

*** Highlights denote the ground truth of each *M* sample.
